# A parametrized two-domain thermodynamic model explains diverse mutational effects on protein allostery

**DOI:** 10.1101/2023.08.06.552196

**Authors:** Zhuang Liu, Thomas Gillis, Srivatsan Raman, Qiang Cui

## Abstract

New experimental findings continue to challenge our understanding of protein allostery. Recent deep mutational scanning study showed that allosteric hotspots in the tetracycline repressor (TetR) and its homologous transcriptional factors are broadly distributed rather than spanning well-defined structural pathways as often assumed. Moreover, hotspot mutation-induced allostery loss was rescued by distributed additional mutations in a degenerate fashion. Here, we develop a two-domain thermodynamic model for TetR, which readily rationalizes these intriguing observations. The model accurately captures the in vivo activities of various mutants with changes in physically transparent parameters, allowing the data-based quantification of mutational effects using statistical inference. Our analysis reveals the intrinsic connection of intra- and inter-domain properties for allosteric regulation and illustrate epistatic interactions that are consistent with structural features of the protein. The insights gained from this study into the nature of two-domain allostery are expected to have broader implications for other multidomain allosteric proteins.

## Introduction

Allostery, a fundamental regulatory mechanism of biomolecular functions, is prevalent in life processes Bacon (1965); Koshland Jr et al. (1966); Changeux and Edelstein (2005); Cui and Karplus (2008); Motlagh et al. (2014); Yu and Koshland Jr (2001); Süel et al. (2003); Dokholyan (2016). The long-range signaling of allostery makes it a fascinating phenomenon, in which binding of an effector molecule (ligand) at the allosteric site alters the function of a distal active site Dokholyan (2016); Leander et al. (2020); Peracchi and Mozzarelli (2011); Wodak et al. (2019). Current descriptions of allostery can be largely cast into two categories: one adopts a mechanical view, focusing on the propagation of conformational distortions from the allosteric site to the active site Wang et al. (2020); Lockless and Ranganathan (1999); Daily and Gray (2009); Rodriguez et al. (2010); Lee et al. (2008); Walker et al. (2020); the other emphasizes the thermodynamic aspect of the problem, highlighting the effect of ligand binding on shifting the protein population among pre-existing conformational states characterized by different ligand binding affinities and active site properties (e.g., the classic MWC model) Bacon (1965); Koshland Jr et al. (1966); Changeux and Edelstein (2005); Cui and Karplus (2008); Sevvana et al. (2012); Takeuchi et al. (2019); Marzen et al. (2013). Models based on both perspectives have provided insights into the function of prototypical allosteric systems thanks to decades of combined efforts of experiment, computation and theory Cui and Karplus (2008); Motlagh et al. (2014); Dokholyan (2016); Marzen et al. (2013); Changeux (2012); Guo and Zhou (2016); Schueler-Furman and Wodak (2016); Reichheld et al. (2009); Xu et al. (2003); Nierzwicki et al. (2021); East et al. (2019). Of note, these two perspectives of allostery are complementary rather than contradictory to each other Liu and Nussinov (2016). The conformational coupling between spatially distant functional sites (allosteric site and active site) plays a vital role in regulating allosteric function, allowing for the transmission of signals from one site to the other Zhang et al. (2020). While the mechanical view primarily seeks to identify the structural basis for signal transduction, it is implicitly assumed within the population shift perspective, which offers a comprehensive and quantitative description of allostery Szabo and Karplus (1972); Viappiani et al. (2014); Henry et al. (2020); Eaton (2022).

In recent years, a thermodynamic model referred to as the ensemble allosteric model (EAM) has been applied to conceptualize protein allostery in terms of intra- and inter-domain properties, with the latter explicitly quantifying the energetic coupling between distant functional sites Motlagh et al. (2014); Wodak et al. (2019); Hilser et al. (2012). This framework is consistent with the observation that allosteric proteins often partition different activities into distinct domains, such as the ligand- and DNA-binding domains in transcription factors and the effector-binding and catalytic domains in enzymes Ramos et al. (2005); Tzeng and Kalodimos (2012); Velyvis et al. (2007); Lipscomb and Kantrowitz (2012). Such thermodynamic approach finds broad applicability across proteins in general, as it has been proposed that all proteins are potentially allosteric Zhang et al. (2020); Gunasekaran et al. (2004). Consequently, this raises intriguing questions about the nature of allostery. For instance, do intrinsic connections exist between the intra- and inter-domain properties of a protein, given the highly cooperative nature of allosteric networks? What roles do sequence and structure play in synergistically determining these properties? Furthermore, what are the parameters within the model that are most essential to the accurate description of realistic allosteric systems, especially prediction of activity upon multiple mutations? To answer these questions and deepen our understanding of allosteric regulation, it is essential to parameterize and test the thermodynamic model using comprehensive mutational data, a topic that still requires further exploration Leander et al. (2020, 2022).

A critical test of any thermodynamic model is whether it can explain the effect of mutations on allosteric signaling. Deep mutational scanning (DMS) analysis has emerged as a powerful function-centric approach over the past decade, which measures the impact of all possible single mutations Fowler et al. (2010); Fowler and Fields (2014); Sarkisyan et al. (2016); Flynn et al. (2019); Starr et al. (2020); Huss et al. (2021). The methodology provides an unbiased way of identifying critical residues for protein allostery and generates extensive data for validating existing computational and theoretical models. Along this line, recent DMS study of four homologous bacterial transcription factors (TFs) in the TetR family (TetR, TtgR, MphR and RolR) revealed that the residues critical for allosteric signaling (hotspots) in these TFs are broadly distributed with no apparent structural link to either the allosteric or the active site Leander et al. (2022); Tack et al. (2021); Faure et al. (2022); Jones et al. (2020); McCormick et al. (2021). This contrasts the commonly held view that hotspot residues tend to form well-defined pathways linking the two sites Süel et al. (2003); Ota and Agard (2005); Strickland et al. (2008); Reynolds et al. (2011); Amor et al. (2016); the observations also hinted at common molecular rules of allostery in the TetR family of TFs Cuthbertson and Nodwell (2013); Fukami-Kobayashi et al. (2003). Moreover, systematic analysis of higher order TetR mutants in the background of five noninducible (“dead”) mutants revealed remarkable functional plasticity in allosteric regulation Leander et al. (2020). Specifically, the loss of inducibility due to the mutations of allostery hotspots could be rescued (restored wildtype-like inducibility) by additional mutations, and different dead mutations exhibit varying numbers of rescuing mutations, which are usually distal and lacking any obvious structural rationale.

While the identification of broadly distributed dead and rescuing mutations in these studies is exciting, such qualitative activity characterization of mutants (inducible and noninducible at a given ligand concentration) prevents a deeper mechanistic understanding of the observation. In fact, mutations exert a graded impact on the allosteric signaling of the TFs; i.e., the expression level of the regulated gene varies among both inducible and noninducible mutants in a continuous, ligand-concentration dependent manner. The nuanced mutational effect on allosteric regulation is linked apparently to the equilibria among different conformational and binding states of the mutant TF, which are determined by the allosteric parameters (intra- and inter-domain properties). Therefore, mapping the mutants onto the parameter space of a biophysical model by exploiting additional experimental data (see below) is crucial for elucidating the observed allosteric phenomena in a comprehensive and physically transparent manner.

In this study, we develop a two-domain statistical thermodynamic model for TetR, in which the protein is generically divided into ligand- and DNA-binding domains (LBD and DBD). Our model incorporates three essential biophysical parameters that capture the intra- and inter-domain properties of the protein, as elaborated in the subsequent section. The model readily rationalizes the myriad ways that mutations can perturb allostery as observed in the aforementioned DMS measurements, by revealing that mutations perturbing intra- and inter-domain properties can lead to similar TF inducibilities at a single ligand concentration. Moreover, the model elucidates the distinct influences of these parameters on the complete induction curve, serving as a diagnostic tool for dissecting the intricate allosteric effects of mutations. We validate the model by accurately describing the induction curves of a comprehensive set of TetR mutants, thereby enabling the quantification of mutational effects and epistatic interactions Daber et al. (2011); Chure et al. (2019). The insights from this combined theoretical and experimental investigation of TetR allostery are expected to generally apply to other two-domain allosteric systems, such as transcription factors like catabolite activator protein (CAP) Tzeng and Kalodimos (2012), receptors like pentameric ligand-gated ion channels (pLGICs) Sauguet et al. (2015); Hu et al. (2020), and allosteric enzymes like aspartate transcarbamoylase (ATCase) Lipscomb and Kantrowitz (2012); Velyvis et al. (2007).

## Results

### Overview of the two-domain thermodynamic model of allostery

As shown in ***Figure 1A and B***, TetR can be generically divided into a LBD and a DBD, disregarding its homodimeric nature Leander et al. (2020); Takeuchi et al. (2019); Reichheld et al. (2009); Leander et al. (2022); Yuan et al. (2022); Scholz et al. (2004). In the same vein of the classic allostery models Bacon (1965); Koshland Jr et al. (1966), each domain features two (the relaxed/inactive and tense/active) conformations that differ in free energies and binding affinities. For the simplicity and mechanistic clarity of the model, we assume negligible binding affinity of the LBD to the lig- and and the DBD to DNA in their inactive conformations and consider competent binding only for the active ones. Each domain must overcome a free energy increase to transition from the inactive to the active conformation (ɛ_L_ for LBD and ɛ_D_ for DBD). Importantly, the two domains are allosterically coupled, in that there is a free energy penalty γ when both domains adopt the active conformations simultaneously. For example, when the ligand binds to the LBD, selecting its active conformation, it discourages the active conformation of the DBD and therefore DNA binding. This anti-cooperativity establishes the foundation for allosteric regulation in TetR.

**Figure 1.**
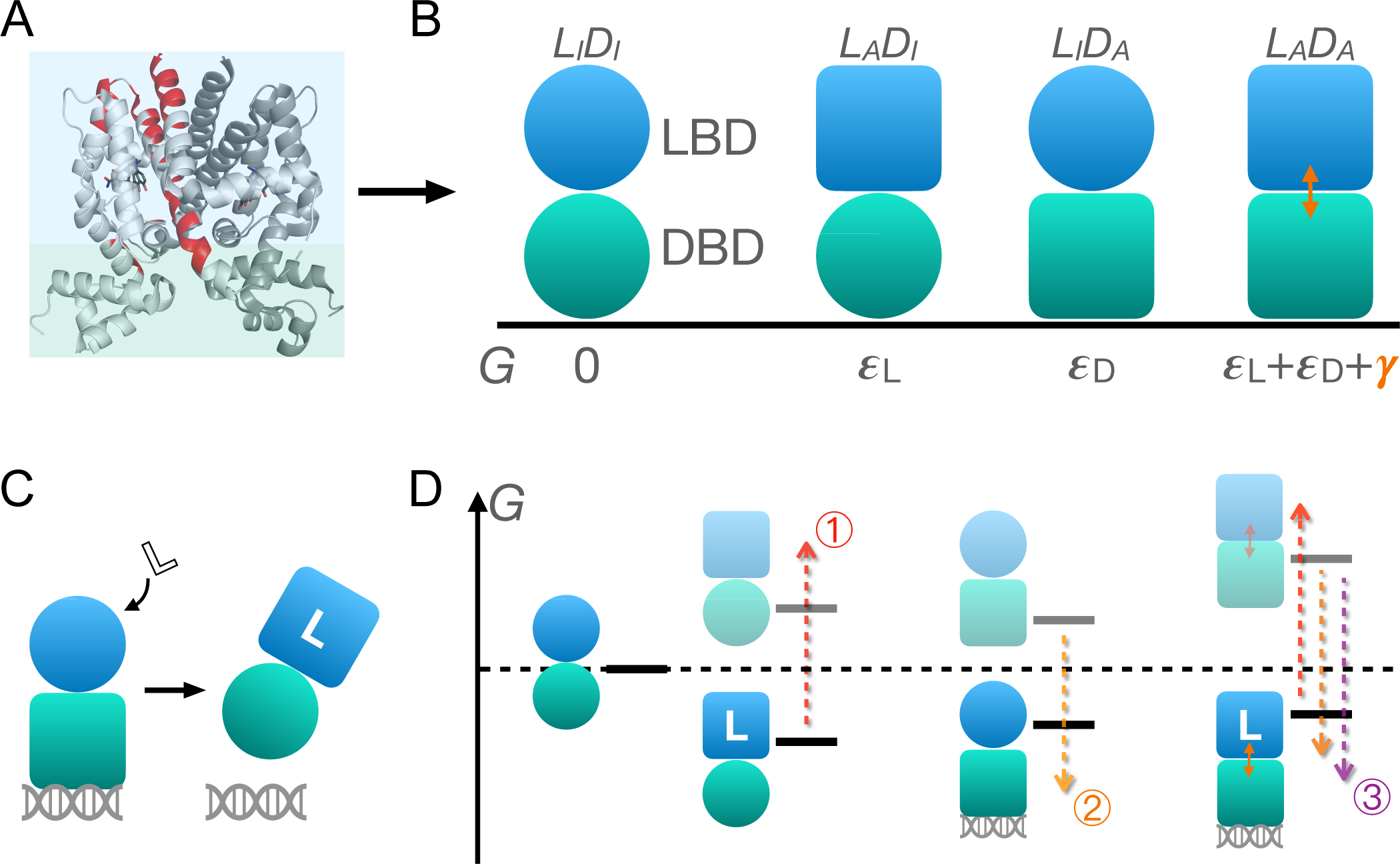
Schematic illustration of the two-domain statistical thermodynamic model of TetR allostery. (A) The crystal structure of TetR(B) in complex with minocycline and magnesium (PDB code: 4AC0). The red residues are the hotspots identified in the DMS study Leander et al. (2020). (B) Four possible conformations of a two-domain TetR molecule with their corresponding free energies (*G*). *G* of the L_I_ D_I_ state is set to 0. Blue/green circle (square) denotes the inactive (active) state of LBD/DBD. (C) A simple repression scheme of TetR function. Binding of the ligand (inducer) favors the inactive state of DBD in TetR, which then releases the DNA operator and enables the transcription of the downstream gene. (D) Schematic free energy diagram of the possible binding states of TetR at fixed ligand and operator concentrations. Red, orange and purple arrows show how a mutation can disrupt allostery by 1. increasing ɛ_L_; 2. decreasing ɛ_D_ and 3. decreasing γ. The L-L_A_ D_A_ state is not explicitly shown in the last column as the doubly-bound L-L_A_ D_A_ -D state is expected to have a lower free energy. Note that mutations that change the binding affinities of the active LBD/DBD to ligand/operator are not discussed here as we focus on the intrinsic allosteric properties of the TF itself.

**Table 1.** Distances to the DNA operator and ligand of the 21 residues under mutational study.

| Residue number | Distance to DNA operator (Å) | Distance to ligand (Å) |
| --- | --- | --- |
| 26 | 7.3 | 24.7 |
| 32 | 12.6 | 30.4 |
| 42 | 8.1 | 25.0 |
| 44 | 9.4 | 21.6 |
| 47 | 7.7 | 21.9 |
| 49 | 11.4 | 17.8 |
| 53 | 17.0 | 12.1 |
| 57 | 22.5 | 7.0 |
| 76 | 45.7 | 17.9 |
| 98 | 19.9 | 15.6 |
| 102 | 18.1 | 14.2 |
| 105 | 24.8 | 7.7 |
| 132 | 39.3 | 16.0 |
| 143 | 29.4 | 16.9 |
| 146 | 25.8 | 17.6 |
| 147 | 26.8 | 16.0 |
| 150 | 23.2 | 19.0 |
| 174 | 34.5 | 14.9 |
| 176 | 38.8 | 19.5 |
| 177 | 35.5 | 17.5 |
| 203 | 55.1 | 28.7 |

**Table 2.** Bayesian inference results with different prior distributions of γ.

| Mutant | $\epsilon_L^1$ | $\epsilon_L^2$ | $\gamma^1$ | $\gamma^2$ |
| --- | --- | --- | --- | --- |
| WT | 6.61<sup>6.73</sup><sub>6.51</sub> | 6.62<sup>6.73</sup><sub>6.51</sub> | 5.29<sup>5.38</sup><sub>5.20</sub> | 5.29<sup>5.38</sup><sub>5.20</sub> |
| Q32A-E147G | 7.29<sup>7.53</sup><sub>7.02</sub> | 7.27<sup>7.53</sup><sub>7.02</sub> | 2.45<sup>2.52</sup><sub>2.38</sub> | 2.45<sup>2.52</sup><sub>2.38</sub> |
| R49G | 8.62<sup>8.84</sup><sub>8.40</sub> | 8.62<sup>8.85</sup><sub>8.42</sub> | 1.66<sup>1.72</sup><sub>1.60</sub> | 1.65<sup>1.71</sup><sub>1.59</sub> |
| D53H | 5.46<sup>5.80</sup><sub>5.09</sub> | 5.46<sup>5.79</sup><sub>5.05</sub> | 1.62<sup>1.66</sup><sub>1.57</sub> | 1.62<sup>1.67</sup><sub>1.57</sub> |
| P105M | 7.23<sup>7.77</sup><sub>6.70</sub> | 7.24<sup>7.76</sup><sub>6.59</sub> | 0.92<sup>0.98</sup><sub>0.86</sub> | 0.92<sup>0.98</sup><sub>0.86</sub> |
| Y132A | 2.35<sup>2.55</sup><sub>2.15</sub> | 2.36<sup>2.55</sup><sub>2.17</sub> | 6.23<sup>6.54</sup><sub>5.97</sub> | 6.22<sup>6.55</sup><sub>5.96</sub> |
| G143M | 6.92<sup>7.18</sup><sub>6.67</sub> | 6.91<sup>7.15</sup><sub>6.68</sub> | 1.52<sup>1.56</sup><sub>1.47</sub> | 1.52<sup>1.56</sup><sub>1.47</sub> |
| E150Y | 6.39<sup>6.73</sup><sub>6.01</sub> | 6.38<sup>6.73</sup><sub>5.99</sub> | 1.72<sup>1.79</sup><sub>1.66</sub> | 1.72<sup>1.78</sup><sub>1.66</sub> |
| PIF | 11.05<sup>11.17</sup><sub>10.92</sub> | 11.05<sup>11.18</sup><sub>10.90</sub> | 1.54<sup>1.59</sup><sub>1.49</sub> | 1.54<sup>1.59</sup><sub>1.49</sub> |
| C203V | 4.51<sup>4.74</sup><sub>4.27</sub> | 4.53<sup>4.78</sup><sub>4.27</sub> | 6.95<sup>10.23</sup><sub>5.89</sub> | 7.47<sup>15.84</sup><sub>6.01</sub> |
| G102D-T26A | 7.23<sup>7.36</sup><sub>7.09</sub> | 7.22<sup>7.35</sup><sub>7.11</sub> | 4.31<sup>4.51</sup><sub>4.16</sub> | 4.31<sup>4.52</sup><sub>4.15</sub> |
| G102D-Y42M-I57N | 8.21<sup>8.34</sup><sub>8.09</sub> | 8.21<sup>8.34</sup><sub>8.09</sub> | 2.83<sup>2.89</sup><sub>2.78</sub> | 2.83<sup>2.88</sup><sub>2.78</sub> |
| G102D-K98Q | 5.96<sup>6.36</sup><sub>5.51</sub> | 5.97<sup>6.37</sup><sub>5.55</sub> | 2.14<sup>2.28</sup><sub>2.01</sub> | 2.14<sup>2.29</sup><sub>2.00</sub> |
| G102D-L146A | 5.70<sup>5.88</sup><sub>5.51</sub> | 5.70<sup>5.88</sup><sub>5.52</sub> | 5.43<sup>6.05</sup><sub>5.03</sub> | 5.42<sup>6.05</sup><sub>5.03</sub> |
| G102D-HQQ | 7.56<sup>7.83</sup><sub>7.30</sub> | 7.57<sup>7.86</sup><sub>7.23</sub> | 1.54<sup>1.61</sup><sub>1.48</sub> | 1.54<sup>1.60</sup><sub>1.48</sub> |
| Y132A-G102D-T26A | 9.55<sup>10.25</sup><sub>8.77</sub> | 9.67<sup>10.51</sup><sub>8.92</sub> | -2.96<sup>-2.58</sup><sub>-3.39</sub> | -3.02<sup>-2.66</sup><sub>-3.47</sub> |
| Y132A-R49G | 5.77<sup>5.98</sup><sub>5.56</sub> | 5.78<sup>5.98</sup><sub>5.58</sub> | 3.11<sup>3.16</sup><sub>3.05</sub> | 3.11<sup>3.16</sup><sub>3.05</sub> |
| Y132A-PIF | 8.71<sup>8.93</sup><sub>8.48</sub> | 8.70<sup>8.93</sup><sub>8.47</sub> | 3.69<sup>3.84</sup><sub>3.57</sub> | 3.69<sup>3.83</sup><sub>3.57</sub> |
| Y132A-C203V | 4.14<sup>4.32</sup><sub>3.94</sub> | 4.15<sup>4.34</sup><sub>3.96</sub> | 7.82<sup>11.12</sup><sub>6.46</sub> | 9.30<sup>16.72</sup><sub>6.76</sub> |
| C203V-R49G | 5.40<sup>5.63</sup><sub>5.14</sub> | 5.39<sup>5.66</sup><sub>5.13</sub> | 2.38<sup>2.43</sup><sub>2.33</sub> | 2.38<sup>2.44</sup><sub>2.33</sub> |
| C203V-D53H | 5.26<sup>5.53</sup><sub>4.98</sub> | 5.26<sup>5.51</sup><sub>4.97</sub> | 1.42<sup>1.46</sup><sub>1.39</sub> | 1.43<sup>1.46</sup><sub>1.39</sub> |
| C203V-G102D-L146A | 3.13<sup>3.43</sup><sub>2.81</sub> | 3.16<sup>3.49</sup><sub>2.78</sub> | 6.74<sup>9.01</sup><sub>6.00</sub> | 6.99<sup>13.63</sup><sub>6.03</sub> |
| C203V-PIF | 8.30<sup>8.61</sup><sub>7.95</sub> | 8.30<sup>8.65</sup><sub>7.94</sub> | 3.49<sup>3.64</sup><sub>3.36</sub> | 3.49<sup>3.64</sup><sub>3.35</sub> |

Accordingly, there are four possible conformational states of TetR, namely L_I_ D_I_, L_A_ D_I_, L_I_ D_A_ and L_A_ D_A_, with L/D and I/A denoting LBD/DBD and inactive/active conformation, respectively (see ***Figure 1B*** and the upper four states of ***Figure 1D***). Binding of ligand/operator to the active LBD/DBD further lowers the free energy of the corresponding state in a concentration dependent manner following the standard formulation of binding equilibrium (see the lower three states of ***Figure 1D***). The regulatory mechanism of TetR allostery can then be qualitatively explained by the schematics in ***Figure 1C and D***. Without the ligand, wildtype (WT) repressor predominantly binds to the operator sequence, which obstructs the binding of RNA polymerase (RNAP) to the adjacent promoter of the regulated gene (***Figure 2—figure Supplement 1A***). In the presence of ligand (inducer) at a sufficiently high concentration, the ligand-bound L_A_ D_I_ state (L-L_A_ D_I_) has the lowest free energy compared with other possible (DNA-bound) states, thus the repressor predominantly releases the operator upon ligand binding, enabling the expression of downstream genes (***Figure 1C*** and ***Figure 2—figure Supplement 1A***).

**Figure 2.**
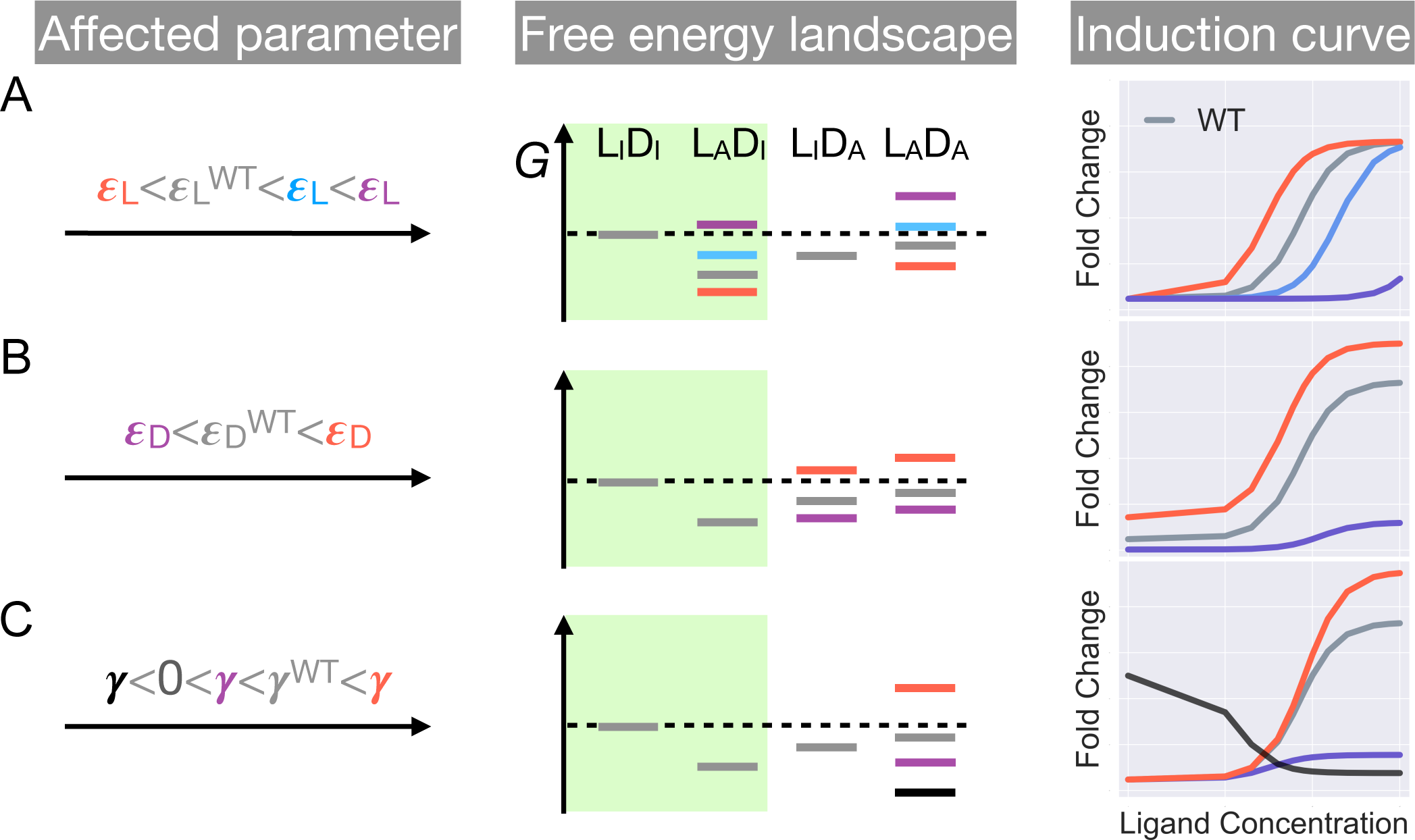
Schematic illustration of the characteristic effects of perturbations in the three biophysical parameters on the free energy landscape and the corresponding induction curves of a two-domain allosteric system. Panels A, B and C illustrate how changing ɛ_L_, ɛ_D_, and γ alone affects the free energy landscape for the binding states shown in ***Figure 1***D and the induction curve. For the black induction curve in (C), the values of ɛ_L_ and ɛ_D_ are also adjusted to aid visualization of the negative monotonicity of the gene expression level (fold change) as a function of ligand concentration. The green shade in the middle column separates the DNA-bound states from the rest. In the free energy landscapes shown in the middle column, ligand- or DNA-binding is always assumed when the corresponding domain is in the active conformation. **Figure 2—figure supplement 1.** Statistical weights of promoter occupancy states and repressor states. **Figure 2—figure supplement 2.** Equilibria among different conformational and binding states of the repressor. **Figure 2—figure supplement 3.** Extended parametric study of main text ***Equation 1***.

Mutations perturb the three biophysical parameters (ɛ_D_, ɛ_L_ and γ) and hence the free energy landscape (***Figure 1***D), leading to changes in TetR function. The three limiting scenarios are depicted in ***Figure 1***D: 1. when a mutation significantly increases ɛ_L_, it raises the free energy of the L-L_A_ D_I_ state relative to the DNA-bound state L_I_ D_A_ -D, which results in a dead (noninducible) phenotype; 2. when a mutation substantially decreases ɛ_D_, it raises the free energy of the L-L_A_ D_I_ state relative to both DNA-bound states (L_I_ D_A_ -D and L-L_A_ D_A_ -D), also leading to a dead mutant; 3. mutations that decrease γ greatly lower the free energy of the double-bound state (L-L_A_ D_A_ -D) relative to L-L_A_ D_I_, again leading to the suppression of induction. Therefore, these schemes highlight that hotspot mutations can disrupt allostery by perturbing either intra- (ɛ_D_, ɛ_L_) or inter-domain (γ) properties. Functional readouts like DMS can help distinguish these mechanistic differences at the biophysical level. Notably, mutations mainly perturbing ɛ_D_ or ɛ_L_ do not have to lie on the structural pathways linking the allosteric and active sites, which explains the broad hotspot distributions observed in the DMS measurement Leander et al. (2020); Reichheld et al. (2009); Leander et al. (2022); Scholz et al. (2004). In general, a dead mutation is likely of mixed nature, as long as its effect on the free energy landscape promotes the dominance of the DNA-bound states. Likewise, rescuing mutations may restore WT-like inducibility by modifying intra- and inter-domain energetics in various ways, as far as the dominance of the L-L_A_ D_I_ state is re-established, rationalizing the broad rescuing mutation distributions. Naturally, the rescuability of a dead mutation depends on which biophysical parameters it perturbs and by how much. At a qualitative level, the distributed nature of dead and rescuing mutations observed in the DMS measurement emerges naturally from the two-domain model Leander et al. (2020).

In summary, the interplay of intra- and inter-domain properties that govern the free energy landscape of TetR’s conformational and binding states, as elucidated by the two-domain model, aligns well with the unexpected DMS results on a qualitative level. To gain deeper insight into TetR allostery and the model itself, we aim to establish a quantitative framework with the model that accurately captures mutant induction curves in the next section.

### System-level ramifications of the two-domain model

The discussions in the previous section and ***Figure 1*** are primarily meant to give an intuitive and qualitative understanding of the two-domain model and rationalization of recent DMS measurements Leander et al. (2020, 2022). In this section, we further establish a quantitative connection between the model and the induction curves of TetR variants, leveraging the recent success of linking sequence-level perturbations to system-level responses Daber et al. (2011); Chure et al. (2019); Garcia and Phillips (2011); Brewster et al. (2014); Weinert et al. (2014); Rydenfelt et al. (2014); Razo-Mejia et al. (2017). An induction curve describes the in vivo expression level of the TetR-regulated gene as a function of inducer concentration.

As described in the last section, the intra-domain parameters (ɛ_D_, ɛ_L_) and inter-domain parameter (γ) of the two-domain model collectively determine the free energy landscape of TetR (***Figure 1***). Consequently, a degenerate relationship arises between combinations of parameter values and the induction level at a specific ligand concentration as measured in the DMS study. To tease apart the allosteric effects of different parameters, we aim to formulate their connection to the full induction curve, which is characterized by (1) the expression level of the TF-regulated gene without ligand (leakiness), (2) gene expression level at saturating ligand concentration (saturation) and (3) the ligand concentration required for half-maximal expression (EC_50_). As revealed in previous studies, these induction curve properties encode information for the key parameters of an allosteric regulation system, for example, the TF binding affinity to ligand and operator, equilibrium between different conformational states of the TF and the abundance of various essential molecules within the system. The establishment of a quantitative connection between the MWC model and the induction curve has yielded valuable insights into several allosteric systems Eaton (2022); Daber et al. (2011); Chure et al. (2019). However, the analogous aspects within a multi-domain thermodynamic model for allostery require further investigation.

Inspired by the pioneering works on transcription with MWC models Daber et al. (2011); Chure et al. (2019), we derived a quantitative relation between the gene expression level in cells and the three biophysical parameters of the two-domain model (***Equation 1***). Briefly, the ratio of the expression level of a TF-regulated gene to that of an unregulated gene, termed fold change (F C) and bound between 0 and 1, is used to quantify gene expression. The gene expression level on the other hand, is assumed to be proportional to the probability that a promoter in the system is bound by RNA polymerase, p(RNAP). Thus, F C is evaluated as the ratio of p(RNAP) in the presence of the TF to that in the absence of the TF. Intuitively, p(RNAP) can be calculated based on the equilibria among different binding and conformational states of the TF (***Figure 2***—figure Supplement 1 and ***Figure 2***—figure Supplement 2), which are determined by the allosteric properties (ɛ_D_, ɛ_L_ and γ) and several other parameters. Therefore, F C can be expressed as a function of these parameters, which under the assumption that intra- and inter-domain energetics adopt typical values of several *k*_B_ *T*, is simplified to ***Equation 1***.

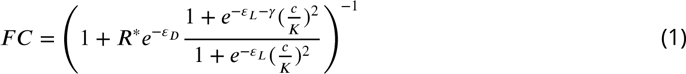

Here, *R*^∗^ is the rescaled TF copy number in the cell; *c* is ligand concentration, and *K* is the dissociation constant of ligand to the repressor with an active LBD. As we focus on understanding how sequence level perturbations are manifested in the allosteric properties of the protein (depicted by ɛ_D_, ɛ_L_ and γ), the residues we choose for mutational study here are mostly not in direct contact with either the ligand or the operator (see supplementary file for more discussions). Thus, *K* and *R*^∗^, which is determined by the TF copy number in the cell and the affinity of the operator to the repressor with an active DBD, are taken to be constants across mutations in all subsequent discussions Chure et al. (2019). Detailed derivations of ***Equation 1*** are provided in the supplementary file.

Despite the considerable degeneracy in the activity (inducible and noninducible) within the parameter space of the two-domain model (***Figure 1***D), ***Equation 1*** demonstrates that mutations affecting distinct biophysical parameters can be discerned based on their characteristic effects on the induction curve.

First, as ɛ_L_ and γ have no effect on the leakiness of the induction curve, only mutations that modify ɛ_D_ can lead to its changes (***Figure 2*** and Supplementary file for additional discussions). In addition, these mutations also change the level of saturation (F C value where the induction curve plateaus at large *c*). Thus, dead mutations that disrupt allostery by decreasing ɛ_D_ alone will uniquely feature a noticeably lower leakiness and a lower level of saturation compared with the WT (***Figure 1***D and ***Figure 2***, see also Supplementary file and ***Figure 2***—figure Supplement 3 for more discussions). Second, mutations that solely perturb the other intra-domain property ɛ_L_, a crucial determinant of TetR’s ligand detection limit, primarily shift the EC_50_ of the induction curve (see ***Equation 1***). As ɛ_L_ increases/decreases from the WT value, the induction curve is right/left shifted with leakiness and the level of saturation remaining unchanged (see the blue and red curves of ***Figure 2***A). However, when ɛ_L_ further increases, the induction curve loses the sigmoidal shape, with its sharply varying tail region being the characteristic of a dead mutation that disrupts allostery mainly by increasing ɛ_L_ (purple curve of ***Figure 2***A, see Supplementary file and ***Figure 2***—figure Supplement 3 for more discussions).

Finally, in the high concentration limit, ***Equation 1*** converges to a constant value (see Equation 2).

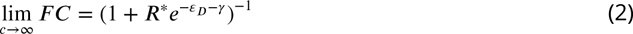

Hence, mutations affecting γ alone will tune the saturation of a sigmoidal induction curve, which increases/decreases as γ increases/decreases (see red and purple curves of ***Figure 2***C). Therefore, in contrast to the two aforementioned scenarios, dead mutations that disrupt allostery through decreasing γ alone will feature a full sigmoidal induction curve with a low level of saturation and unchanged leakiness compared with the WT (with γ > 0, see the purple curve of ***Figure 2***C). Furthermore, as γ dictates the inter-domain cooperativity, it thereby controls the sign of the monotonicity of F C as a function of *c* (see Supplementary file for detailed proof). Specifically, when γ/0, there is negative/positive cooperativity between ligand and operator binding, and F C increases/decreases monotonically with *c* (see ***Figure 2***C). When γ = 0, however, the bindings of ligand and operator become independent of each other, and *c* no longer affects F C (***Figure 2***—figure Supplement 3).

The distinctive roles of the three biophysical parameter on the induction curve as stipulated in ***Equation 1*** could be understood in an intuitive manner as well. First, the value of ɛ_D_ controls the intrinsic strength of binding of TetR to the operator, or the intrinsic difficulty for ligand to induce their separation. Therefore, it controls how tightly the downstream gene is regulated by TetR without ligands (reflected in leakiness) and affects the performance limit of ligands (reflected in saturation). Second, the value of ɛ_L_ controls how favorable ligand binding is in free energy. When ɛ_L_ increases, the binding of ligand at low concentrations become unfavorable, where the ligands cannot effectively bind to TetR to induce its separation from the operator. Therefore, the fold-change as a function of ligand concentration only starts to noticeably increase at higher ligand concentrations, resulting in larger EC_50_. Third, as discussed above, γ controls the level of anti-cooperativity between the ligand and operator binding of TetR, which is the basis of its allosteric regulation. In other words, γ controls how strongly ligand binding is incompatible with operator binding for TetR, hence it controls the performance limit of ligand (reflected in saturation).

Having identified the distinctive impacts of the different allosteric parameters of the two-domain model on the induction curve, as outlined in ***Equation 1***, we next apply the model to analyze actual experimental induction curves of TetR mutants. The analysis enables us to uncover the underlying biophysical factors contributing to diverse mutational effects. Additionally, it allows us to evaluate the model’s validity in capturing TetR allostery by assessing its accuracy in reproducing the experimental data.

### Extensive induction curves fitting of TetR mutants

With the diagnostic tools established, we mapped mutants to the parameter space of the two-domain model through fitting of their induction curves. We choose 15 mutants for analysis in this section, which contain mutations that span different regions in the sequence and structure of TetR (***Figure 3***—figure Supplement 1 and ***Figure 3***—figure Supplement 9). Five of the 15 mutants consist of a dead mutation G102D and one of its rescuing mutations (see the third row of ***Figure 3***A), while the other 10 contains the WT, 8 single mutants and a double mutant Q32A-E147G (see the first two rows of ***Figure 3***A and Supplementary file section 7 for more discussions). In all cases, fitting of an induction curve is divided into two steps; first, at *c* = 0, ***Equation 1*** can be rearranged to give which enables the direct calculation of ɛ_D_ from the leakiness of the induction curve; second, ɛ_L_ and γ are inferred based on the remaining induction data using the method of Bayesian inference Chure et al. (2019). Briefly, given ɛ_D_ calculated in the first step, we select a set of ɛ_L_ and γ values from physical ranges of the parameters, based on the probability of observing the induction curve with the parameter values using Monte Carlo sampling. The medians of the selected sets of ɛ_L_ and γ values, which are known as posterior distributions, are reported as the inferred parameter values, and error bars show the 95% credible regions of the posterior distributions (***Figure 3***B and ***Figure 4***—figure Supplement 2B). All details are provided in the “*Materials and methods*” section and Supplementary file.

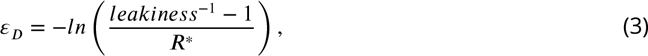

**Figure 3.**
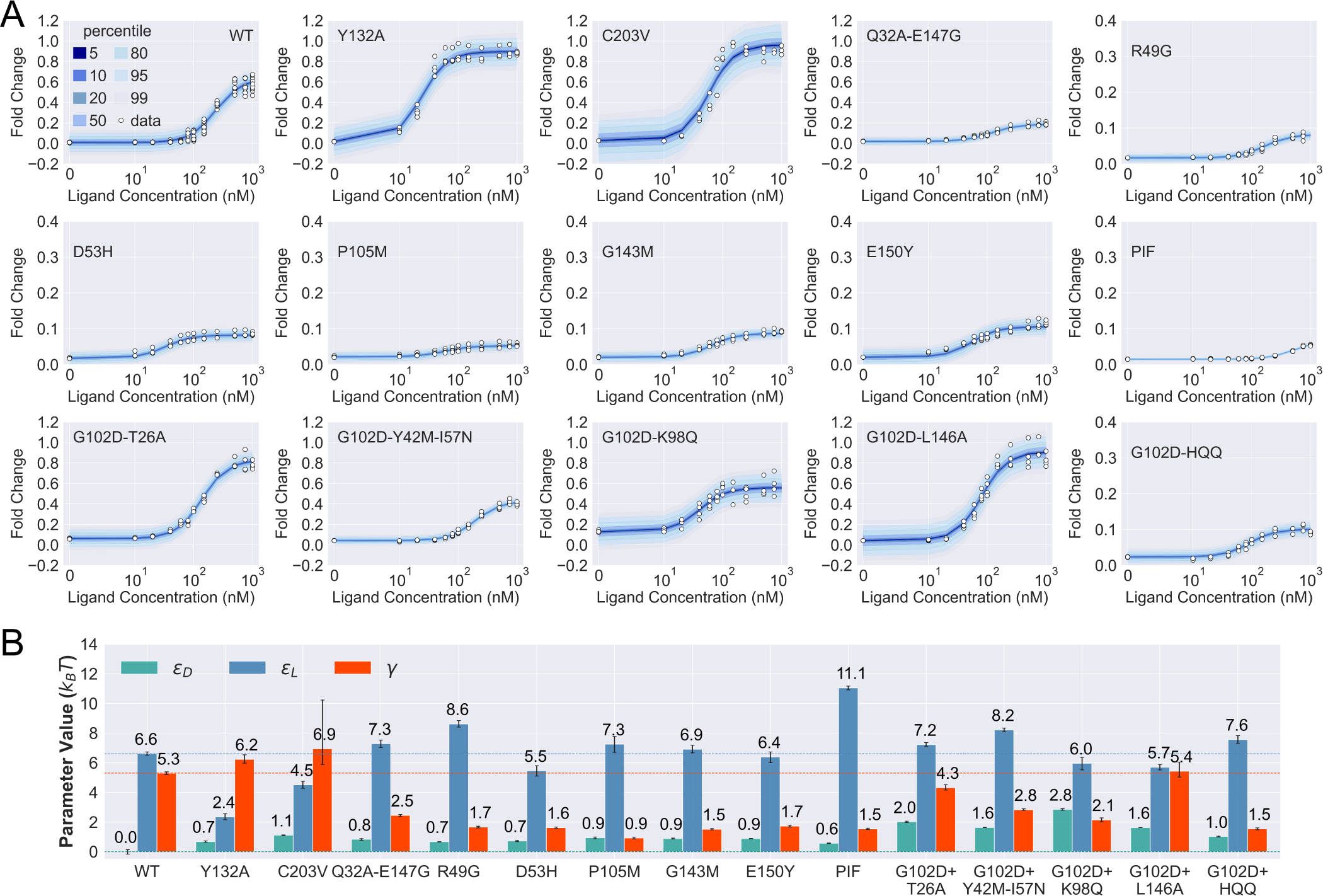
Induction data of 15 TetR mutants and the corresponding parameter estimation results. (A) Shaded blue curves in each plot show the percentiles of the simulated fold-change measurements using the inferred posterior parameters of the mutant. The white data points represent the corresponding experimental induction measurement of 4 or more biological replicates (three replicates for C203V and G102D-HQQ). (B) The inferred parameter values of the 15 mutants. The error bars of ɛ_L_ and γ represent the 95^*th*^percentile of the Bayesian posterior samples, while the error bar of ɛ_D_ is calculated based on the standard error of the mean (SEM) of the corresponding leakiness measurement. The horizontal lines indicate the WT parameter values for reference. **Figure 3—figure supplement 1.** Sequence and structural distributions of the 21 residues chosen for the mutation analyses in this work. **Figure 3—figure supplement 2**. Prior probability distributions and prior predictive check. **Figure 3—figure supplement 3**. Distributions of the prior predictive parameters and the corresponding posterior distributions. **Figure 3—figure supplement** 4. Distributions of rank statistics of the prior predictive parameters relative to the corresponding posterior samples. **Figure 3—figure supplement** 5. Sensitivity analysis for model parameter inference. **Figure 3—figure supplement** 6. Posterior predictive check of mutant G102D-Y42M-I57N. **Figure 3—figure supplement** 7. Theoretical induction curves of R49G, D53H, P105M and G143M with WT γ value. **Figure 3—figure supplement** 8. Sorting scheme to identify dead variants. **Figure 3—figure supplement** 9. Distances to the DNA operator and ligand of the 21 residues under mutational study. **Figure 3—figure supplement** 10. Bayesian inference results with different prior distributions of γ.

**Figure 4.**
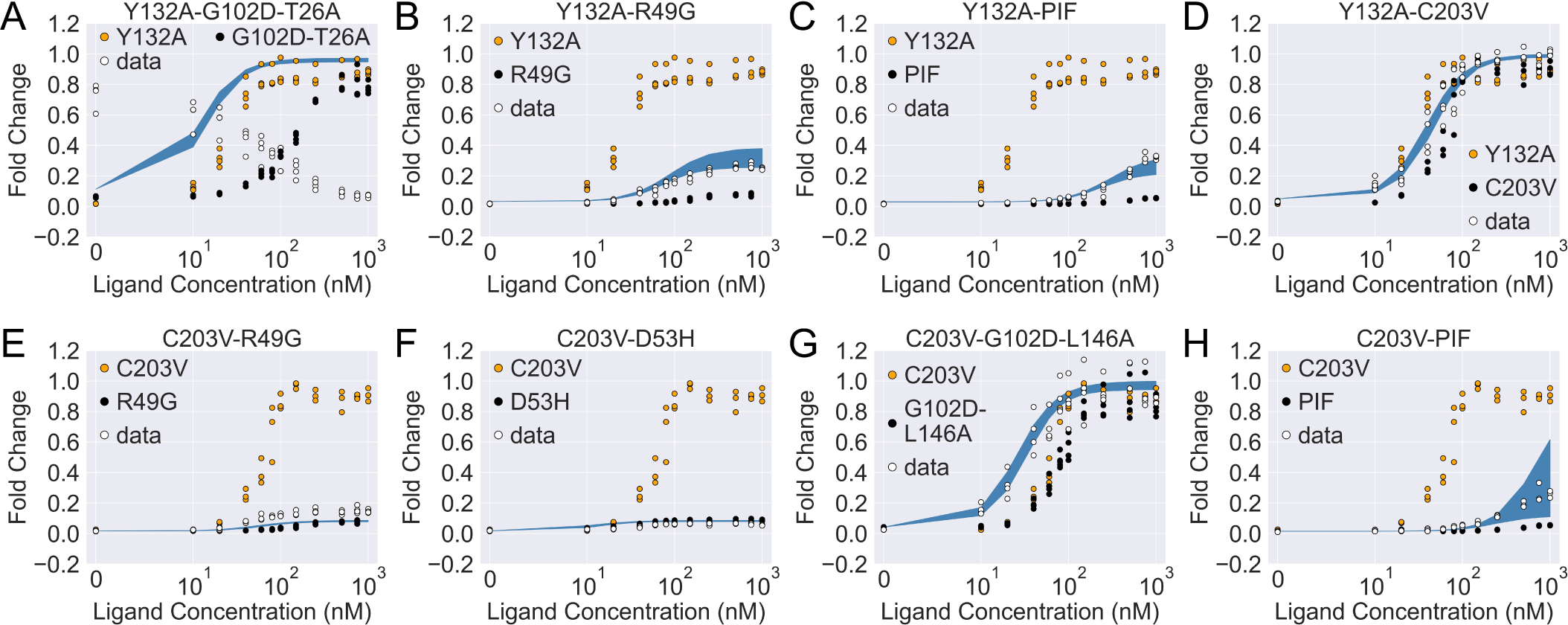
The induction curves for the eight combined TetR mutants from experimental measurement and prediction from the modified additive model. In each plot, black, orange and white points represent the experimental data for mutant 1, mutant 2 and the combined mutant (named mutant 1-mutant 2), specified in the legend and title. The blue band show the 95^*th*^percentile of the indu_{_ ction curve prediction from th_}_ e modified additive model. The modification to the basic additive model in each plot is specified by the 6 weights {1,ɛ_D_ ^, α^2,ɛ_D_ ^, α^1,ɛ_L_ ^, α^2,ɛ_L_ ^, α^1,γ ^, α^2,γ (see ***Equation 4***), which are (A) {1, 1, 1, 1, 1, 1}; (B) {1, 1, 0.5, 1, 1, 1}; (C) {1, 1, 0.5, 1, 1, 1}; (D) {1, 1, 0, 1, 1, 1}; (E) {0, 1, 1, 1, 0, 1}; (F) {0, 1, 1, 1, 0, 1}; (G) {0, 1, 1, 1, 1, 1}; (H) {0, 1, 1, 1, 1, 1}. **Figure 4—figure supplement 1.** The induction curves of the eight combined mutants calculated using the basic additive model. **Figure 4—figure supplement 2.** Induction curves of the eight combined mutants and the corresponding parameter estimation results as well as the basic additive model predictions.

As shown in ***Figure 3***A, the fitted curves accurately capture the induction data of all the mutants studied here, enabling the quantification of their biophysical parameters with little uncertainty (see ***Figure 3***B and ***Figure 3***—figure Supplement 10). These results support the general applicability of the two-domain model to describing TetR allostery, as these 15 mutants vary significantly in terms of location of mutations and phenotype. They are one to four mutations away from the WT, and the sites of mutations are distributed across LBD, DBD and the domain interface (***Figure 3***—figure Supplement 1). In terms of the phenotype, they fall into different classes as characterized by the F C value at *c* = 1000 nM (F C^1000^), including dead (F C^1000^ < 0.1), enhanced induction (F C^1000^ > F C^1000^) and neutral activity (the rest).

Besides inspecting the goodness of fit, a closer examination of the induction curves and the Bayesian inference results reveal additional features of the two-domain model of TetR allostery. First, the apparent binding affinity of TetR to the ligand (anhydrotetracycline; aTC) and operator (*tet02*) used in our experiments are estimated to be about 17.4 *k*_B_ *T* and 16.4 *k*_B_ *T*, respectively, based on the parameter values inferred for the WT (see Supplementary file for detailed calculations); these agree well with the range of reported experimental values Scholz et al. (2000); Schubert et al. (2004); Kędracka-Krok and Wasylewski (1999); Kamionka et al. (2004); Bolintineanu et al. (2014); Normanno et al. (2015). Second, as shown in ***Figure 3***B, the perturbations in ɛ_D_ are generally small in magnitude (< 1 *k*_B_ *T*) among the dead mutants, while substantially larger perturbations (up to ∼ 5 *k*_B_ *T*) are observed for ɛ_L_ and γ. The latter observation is in line with the predictions of the model (see the previous section) that two types of qualitatively different induction curves are expected for partially dead mutants (see the first and second rows of ***Figure 3***A). Specifically, the induction curve of P176N-I174K-F177S (PIF) loses the sigmoidal shape, with its sharply varying tail region suggesting a significant increase of ɛ_L_. This is confirmed by the parameter fitting result, which shows that the triple mutations of PIF, located in the core region of LBD (Supplementary file and ***Figure 3***—figure Supplement 1), lead to the largest increase of ɛ_L_ among all mutants (4.5 *k*_B_ *T* higher than the WT, see ***Figure 3***B). The induction curves of the other four dead mutants (R49G, D53H, P105M and G143M), however, maintain the sigmoidal shape yet with low levels of saturation. As they all exhibit a higher ɛ_D_ (leakiness) than the WT, the low levels of saturation have to result solely from weakened inter-domain couplings (***Figure 3***). In other words, these four dead mutations, located mostly at the domain interface (***Figure 3***—figure Supplement 1), disrupt allostery primarily through decreasing γ. Indeed, when the γ of these mutants is set to the WT value but keeping their respective ɛ_D_ and ɛ_L_ parameters unchanged, higher F C^1000^ values than the WT are obtained instead (***Figure 3***—figure Supplement 7).

Despite the discussion of two limiting types of changes in the induction curves, it is interesting to observe in ***Figure 3***B that the three biophysical parameters, especially ɛ_L_ and γ, are often perturbed together in the mutants. Indeed, although the three parameters are theoretically independent of each other, at the structural level, it is less likely to have a scenario where a mutation perturbs inter-domain coupling (γ) but leaves intra-domain properties unchanged. Lastly, while quantification of G102D from its flat induction curve incurs large uncertainties, (***Figure 2***—figure Supplement 3), the diversities of the induction curves and the biophysical parameters of the G102D-rescuing mutants (***Figure 3***B) provide clear support for the qualitative anticipation from the previous section; i.e., rescuing mutations of a dead mutant may restore inducibility through different combinations of tuning ɛ_D_, ɛ_L_ and γ. For example, while Y42M-I57N and K98Q both rescue G102D to achieve similar F C^1000^, they exert distinct influences on ɛ_D_ and ɛ_L_, as reflected in the variations of leakiness and EC_50_ between the corresponding induction curves. In another case, while the induction curves of G102D-T26A and G102D-L146A show comparable values of saturation and leakiness, their different EC_50_ values suggest the different effects of the two rescuing mutations on ɛ_L_, further quantified by the inferred parameter values. The diverse mechanistic origins of the rescuing mutations revealed here provide a rational basis for the broad distributions of such mutations. Integrating such thermodynamic analysis with structural and dynamic assessment of allosteric proteins for efficient and quantitative rescuing mutation design could present an interesting avenue for future research, particularly in the context of biomedical applications Pan et al. (2005); Liu and Nussinov (2008).

In summary, ***Equation 1*** accurately captures the induction curves of a comprehensive set of TetR mutants using a minimum set of parameters with realistic values (see the section “Discussion” and ***Figure 3***—figure Supplement 1). The two-domain model thus provides a quantitative platform for investigating TetR allostery beyond an intuitive rationalization of the DMS results.

### Exploring epistasis between mutations

With the 15 mutants mapped to the parameter space of the two-domain model, we now explore the epistatic interactions between the relevant mutations. To do so, we start by assuming additivity in the perturbation of all three biophysical parameters (see ***Equation 4*** and ***Equation 5***, where *p* represents any one of ɛ_D_, ɛ_L_ and γ).

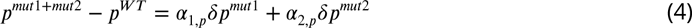

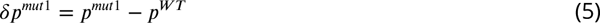

We then evaluate how the induction curves generated by this additive model (α_1,*p*_ = α_2,*p*_ = 1) deviate from the corresponding experimental results, and the magnitude of deviation quantifies the significance of epistasis. Eight mutation combinations were chosen for the analysis, where we pair up C203V and Y132A, the two single mutations that enhance the level of induction relative to the WT, with mutations from different structural regions of TetR (see Supplementary file section 7 for more discussions).

Although predictions from the additive model are qualitatively correct for 5 of the 8 mutation combinations on phenotypic effects, none of them captures the corresponding induction curve accurately (***Figure 4***—figure Supplement 1). This again highlights the parameter space degeneracy of the phenotypes, and that mechanistic specificity is required of a model for making reliable predictions on combined mutants Li and Lehner (2020). For a deeper understanding of epistatic interactions between the mutations queried here, we next directly fit for the three biophysical parameters of the combined mutants using their induction curves, which are then compared with those obtained from the additive model.

As shown in ***Figure 4***—figure Supplement 2A, the induction data of all eight combined mutants are well captured by the fitted curves, which reaffirms the applicability of the two-domain model. Interestingly, despite the discrepancy between the experimental induction curves and those predicted by the additive model, the fitted parameters of the combined mutants are comparable to those from the additive model in many cases (see ***Figure 4***—figure Supplement 2B). In this regard, one noticeable example is the mutant C203V-G102D-L146A, for which the result of direct fitting is close to the additive model in terms of both ɛ_L_ and γ (with a difference of ⩽ 0.5 *k*_B_ *T*), while they differ more in ɛ_D_ (1.6 *k*_B_ *T*). Inspired by such observations, we then seek the minimal modifications to the additive model that can account for epistasis. As C203 is one of the most distant residues from DBD in TetR, we reason that its effect on ɛ_D_ may get dominated by mutations much closer to the DNA binding residues like G102D and L146A (***Figure 3***—figure Supplement 1 and ***Figure 3***—figure Supplement 9). Remarkably, when we quench C203V’s contribution to ɛ_D_ in the additive model of C203V-G102D-L146A (α_C203V,ɛ_ = 0, see ***Equation 4***), the predicted induction curve well recapitulates the experimental data (***Figure 4****G*). In another example, when C203V is combined with the triple mutations PIF, which are much closer to the DBD, the fitted parameters of the combined mutant also align well with the additive model except only for ɛ_D_. Here, quenching C203V’s contribution to ɛ_D_ again leads to good agreement between the additive model and experiment in terms of the induction curve (***Figure 4***H).

The epistatic effects in other C203V-containing combined mutants can be largely accounted for by modifying the additive model following the same physical reasoning as above. For instance, mutation D53H is near the DNA-binding residues and located at the domain interface, which suggests its dominant role in defining the DBD energetics and inter-domain cooperativity when paired up with C203V Scholz et al. (2004). Accordingly, quenching C203V’s effect on both ɛ_D_ and γ in the additive model for mutant C203V-D53H leads to dramatic improvement in the induction curve prediction (***Figure 4***—figure Supplement 1F and ***Figure 4***F). Along this line, success is also observed when accounting for the epistasis between C203V and R49G by the same approach (***Figure 4***E). These examples might also explain why C203V, although being able to enhance the induction of TetR, fails to rescue a range of dead mutations including R49A, D53V, G102D, N129D and G196D Leander et al. (2020). More broadly, these observations together indicate that epistasis in combined mutants (e.g., those containing C203V) can be captured by the additive two-domain model with modifications based on physical reasoning.

The epistasis in combined mutants containing Y132A can be understood in a similar manner. Noticeably, the fitted parameter values of Y132A-C203V agree very well with the additive model for ɛ_D_ and γ (within a difference of 0.4 *k*_B_ *T*), while they differ in ɛ_L_ by 3.8 *k*_B_ *T*. Interestingly, such discrepancy in ɛ_L_ values can be essentially resolved by quenching Y132A’s contribution in the additive model (reduced to 0.4 *k*_B_ *T*). This likely suggests that the effect of Y132A on ɛ_L_ tends to be compromised when combined with other mutations that perturb ɛ_L_. Indeed, for most of the Y132A containing mutants investigated here (Y132A-C203V/PIF/R49G), the accuracy of induction curve prediction by the additive model improves greatly when α_Y132A,ɛ_ is tuned down (***Figure 4***B − D, see Supplementary file for more discussions).

The only exception to this trend is the mutant Y132A-G102D-T26A, for which we observe strong epistasis between Y132A and the dead-rescue mutation pair G102D-T26A. Although both mutations Y132A and G102D-T26A enhance the allosteric response of TetR (***Figure 3***), their combined effects radically change the sign of cooperativity between the two domains (γ), turning the ligand (aTC) from an inducer into a corepressor (***Figure 4***A and ***Figure 4***—figure Supplement 2B). Here, large disparity exists between the additive model and the direct fitting result in all three biophysical parameters, which cannot be explained by simple modifications of the additive model.

## Discussion

Allostery, a major regulatory mechanism in biology, has attracted intense research interest in the past few decades due to its complexity and implications in biomedicine and protein engineering. A central goal is to develop a quantitative understanding of the phenomena through a physical model that can be tested by comprehensive data. Along this line, one of us has in recent years advanced a function-centric approach to studying protein allostery with DMS, which provides a comprehensive and most direct test of our mechanistic understandings. Specifically, we have shown in an unbiased way that allostery hotspots and dead-rescue mutation pairs in four TetR family TFs are distributed across the protein structures Leander et al. (2020, 2022). This highlights that modifying the propagation of conformational distortions between the effector and active sites is not the only way to tune allosteric regulation Leander et al. (2020).

The rich and surprising observations for the TetR family TFs call for a physical understanding of the underlying allostery mechanism, for which we resort to statistical thermodynamic models. Statistical thermodynamic models have played a central role in shaping our understanding of allostery Bacon (1965); Koshland Jr et al. (1966). In particular, Chure et al. have developed the methodology of fitting the MWC model to the induction data of LacI repressors using Bayesian inference Chure et al. (2019). This established the connection between sequence-level perturbations to system-level responses, which enables the exploration of mutational effects on transcription within the framework of biophysical models. However, as pointed out in several previous studies, the MWC model is not consistent with the observation that effector-bound TetR crystal structures are closer to the DNA-bound form compared to the apo crystal structures Motlagh et al. (2014); Reichheld et al. (2009); Hilser et al. (2012). Hence, it’s at least difficult to provide a complete picture of TetR allostery with the MWC model. The ensemble allosteric model (EAM) on the other hand, presents a more flexible and detailed domain-specific view of allostery, which can be applied to more complex allosteric systems (e.g., allostery with intrinsically disordered proteins) Motlagh et al. (2014); Hilser et al. (2012).

Inspired by these pioneering developments and our DMS results, in this work, we propose a two-domain statistical thermodynamic model, the parameter-activity degeneracy of which readily elucidates the distributed allosteric network observed in the DMS result of TetR (***Figure 1***). On the other hand, the functional form derived in ***Equation 1*** establishes the qualitative differences in the impacts of various model parameters on the characteristics of the induction curve (***Figure 2***); for example, dead mutations perturbing inter- and intra-domain properties are predicted to cause induction curves with and without a saturating plateau, which are both observed experimentally (***Figure 3***). Moreover, ***Equation 1*** accurately captures the induction data of a diverse set of TetR mutants (***Figure 3***, ***Figure 3***—figure Supplement 1 and ***Figure 4***—figure Supplement 2) in a quantitative manner, enabling their mapping to the model parameter space with high precision. It’s noted that the homodimeric nature of TetR is ignored in the current two-domain model to minimize the number of parameters, and additional experimental data could necessitate a more complex model for TetR allostery in the future (see Supplementary file section 8 for more discussions).

The success of ***Equation 1*** allows for a quantitative investigation of epistasis between the characterized mutations. In a previous study of another TF, LacI, using the MWC model, no epistasis was observed between mutations of ligand-binding and DNA-binding residues analyzed therein Chure et al. (2019). In the present analysis of TetR, epistasis is observed in all eight queried mutation combinations, which can be largely rationalized through physical reasoning. For example, our results intuitively indicate that the effect of a distant mutation like C203V on DBD energetics and inter-domain coupling tend to be dominated by mutations much closer to these regions (Figure 4 and ***Figure 4***—figure Supplement 1). Such phenomena suggest that these mutations affect allosteric regulation by influencing not only the relative populations of conformations in different ligation states, but also through shifting the dominant conformations themselves Zhang et al. (2020). On the other hand, epistasis involving Y132A is more complex. For instance, we observe that Y132A’s influence on ɛ_L_ is diminished when paired up with other mutations in most cases. Further mechanistic understanding, likely from a biochemical perspective, is required to fully explain this phenomenon. Nonetheless, these observations provide a plausible explanation for why C203V and Y132A, although being able to enhance the induction response of TetR individually, could not rescue a range of dead mutations Leander et al. (2020).

Our results reveal additional insights into the nature of two-domain allostery as well. Although our model assumes no a *priori* correlation between the intra- and inter-domain properties of the TF, we find that they are always modified together by mutations (***Figure 3*** and ***Figure 4***—figure Supplement 2). This immediately points to the interconnectivity of allosteric networks Takeuchi et al. (2019); Reichheld et al. (2009); Scholz et al. (2004). That is, when a residue is involved in the conformational rearrangements induced by both effector and operator binding, it functions as a bridge through which the two events are coupled. Within such a conceptual framework, it is highly unlikely to observe a mutation that changes γ alone without perturbing the intra-domain energetics.

This point is explicitly illustrated in ***Figure 5***, which summarizes the locations of all the characterized 23 mutants in the two-dimensional space of ɛ_L_ and γ. Here, the color of the contour plot encodes the F C^1000^ value calculated for each (ɛ_L_, γ) combination together with the WT ɛ_D_ value, while the color of the specific mutant data points is based on the true F C^1000^ value. Thus, the color of a data point reflects the ɛ_D_ of the corresponding mutant relative to the WT; i.e., a brighter/darker color than the surrounding indicates that the mutant features a higher/lower ɛ_D_ value than the WT. Since the variations of ɛ_D_ among the investigated mutants are modest compared with ɛ_L_ and γ (see ***Figure 3***B), we’ll primarily focus on the latter two in the discussion. Evidently, mutants with the strongest allosteric responses feature small ɛ_L_ and large γ, which correspond to the upper left region of ***Figure 5***. Additional mutations in the background of these mutants lead to weaker allosteric response, moving the mutants to the darker regions of the plot. The interconnected nature of allosteric networks described by the two-domain model, however, dictates that such shifts would take place along the diagonal of the ɛ_L_ -γ plane, which describe mutations that modify intra- and inter-domain properties simultaneously. This feature potentially offers a functional advantage of preventing mutants from regions of low allosteric response (corresponding to upper right and lower left of ***Figure 5***), facilitating efficient adaptation to new effectors during evolution. In the future, it is of interest to examine whether such a negative correlation between ɛ_L_ and γ is a generic feature of two-domain allostery.

**Figure 5.**
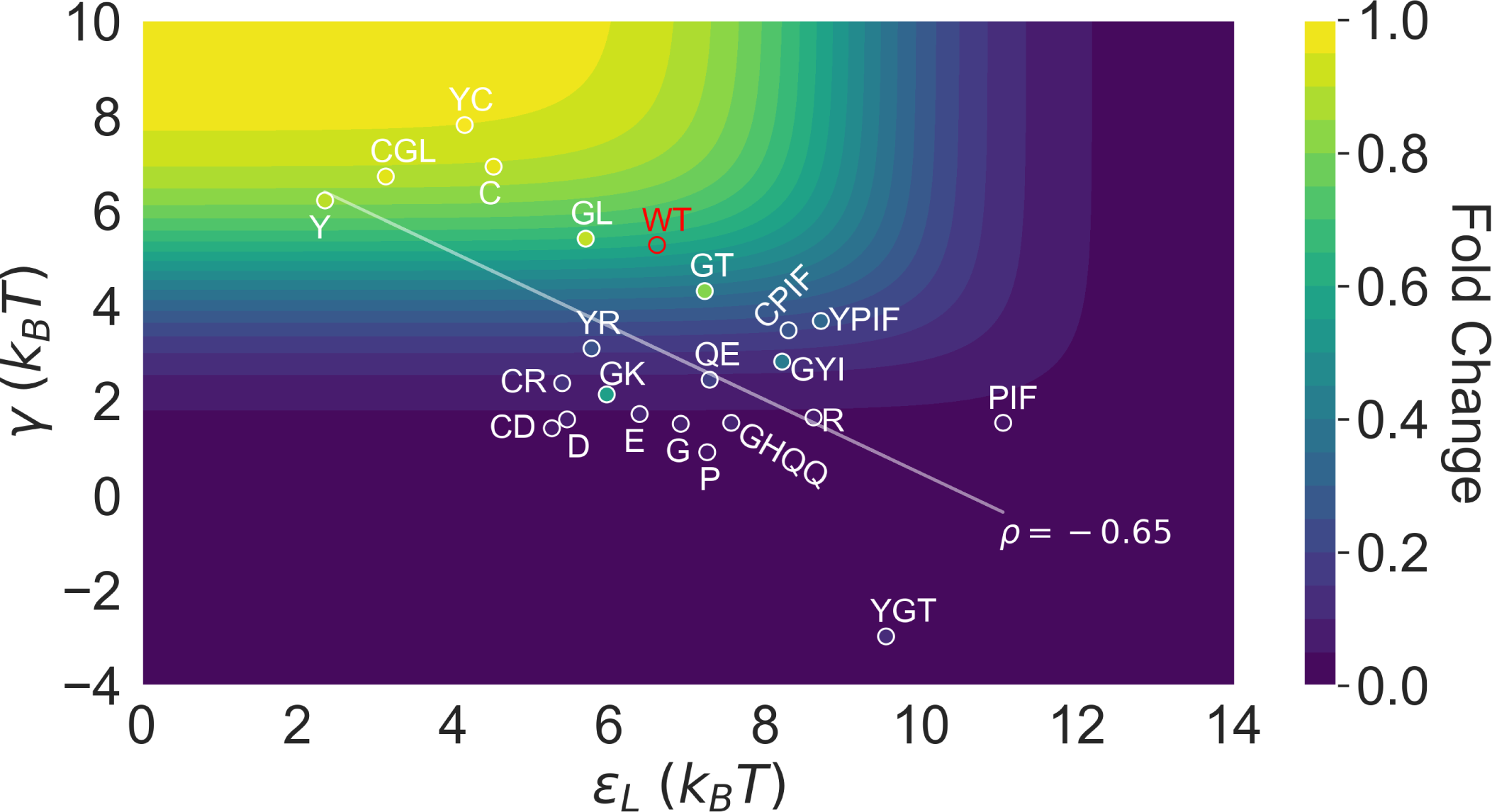
Distribution of the 23 investigated TetR mutants in the parameter space of the two-domain model illustrates the correlation between perturbations in ɛ_L_ and γ. Color of the contour plot encodes the fold change at *c* = 1000 nM calculated for each point in the two-dimensional space of ɛ_L_ and γ with the WT ɛ_D_ value. The color within the data point of each mutant is based on the F C^1000^ calculated with its specific ɛ_D_ value. The notation of each mutant is abbreviated based on the 1-letter codes of the residues that are mutated. The specific mutations corresponding to each letter code from upper left to lower right are C: C203V; Y: Y132A; GL: G102D-L146A; GT: G102D-T26A; R: R49G; D: D53H; GK: G102D-K98Q; E: E150Y; QE: Q32A-E147G; G: G143M; P: P105M; GYI: G102D-Y42M-I57N; GHQQ: G102D-H44F-Q47S-Q76K; PIF: P176N-I174K-F177S. The least-squares regression line between ɛ_L_ and γ is shown with their Pearson correlation coefficient (*ρ*).

Our results establish the two-domain model as a flexible and quantitative platform for investigating TetR allostery, adding to the value of statistical thermodynamic models as advocated by the seminal works on MWC molecules Daber et al. (2011); Chure et al. (2019). We note, however, that the two-domain model is a more generic description of allostery compared with the MWC model.

First, although our model identifies the importance of intra-domain energetics in tuning the free energies of different TF states, it points out the more fundamental role of inter-domain coupling in defining allosteric response (***Figure 2*** and Supplementary file). A large inter-domain coupling is assumed to be obligatory in the MWC model and required to create the two pre-existing conformations of different ligand and operator affinities in the first place. Various experimental studies as well as our own data reveal that the inter-domain coupling of TetR is amenable to tuning, especially by mutations of residues at the domain interface Reichheld et al. (2009); Scholz et al. (2004); Müller et al. (1995); Hecht et al. (1993). In some cases, even single mutations can reverse the sign of cooperativity between LBD and DBD, indicating that the dead mutational effects observed in the DMS experiments and elsewhere could well originate from diminishing γ. These considerations necessitate an explicit treatment of γ in a physical model of TetR allostery.

Second, the MWC model, when applied to two-domain systems, contains at least four biophysical parameters Daber et al. (2011); Chure et al. (2019), is thus more complex than our two-domain model, which requires only three. The simplicity of our model reduces its parameter space degeneracy, enabling high precision in fitting while being able to accurately describe a broad set of induction data. The three parameters also offer an interpretable mechanistic picture of TetR allostery in a physically transparent fashion.

We have focused our experimental and theoretical analyses on the TetR-family TFs, which represent important systems for building allosteric models, due to their broad involvement in many aspects of cell physiology Ramos et al. (2005); Cuthbertson and Nodwell (2013). Along this line, a natural extension of the current study is to use the two-domain model as a platform for comparing allostery in different TetR homologs Leander et al. (2022). For example, coupling the model with systematic dose-response study of hotspot mutations could reveal the roles of different allosteric hotspots (whether they dictate intra- or inter-domain properties or both). The similarity in key structural features among TetR homologs then enables the comparison of the nature of hotspots at similar structural locations. This could potentially lead to novel insights into the different roles that sequence and structure play in defining the allostery of a system, which is a fundamental question to address in rational engineering of allosteric proteins.

Although our two-domain model is inspired by and established with the data of TetR family TFs, the mechanistic and modelling insights gained here should be generally applicable to other allosteric systems sharing the two-domain architecture. This includes other transcription factors like catabolite activator protein (CAP), a homodimeric TF with a cyclic AMP (effector) binding domain and a DNA binding domain Tzeng and Kalodimos (2012); receptors like pentameric ligand-gated ion channels (pLGICs), which allosterically translate the binding of neurotransmitters to their extra-cellular domains into the activation of ionotropic pores located in their transmembrane domains Sauguet et al. (2015); Hu et al. (2020); and allosteric enzymes like aspartate transcarbamoylase (AT-Case), in which binding of effector like ATP to the regulatory domain allosterically alters the catalytic activity of the functional domain Velyvis et al. (2007); Lipscomb and Kantrowitz (2012). Ultimately, we envision that such model provides a minimalist but physically sound framework for integrating different types of experimental data and computational methods Leander et al. (2022); Yuan et al. (2022); Xie et al. (2022); Tonner et al. (2021) for exploring rational engineering and regulation of allostery for biotechnological and biomedical applications.

## Materials and methods

### Library construction

Using a low-copy backbone (SC101 origin of replication) carrying spectinomycin resistance, we constructed a sensor plasmid with TetR(B) (Uniprot P04483). The tet*Rb* gene was driven by a variant of promoter apFAB61 and Bba_J61132 RBS Kosuri et al. (2013). On a second reporter plasmid, superfolder GFP Pedelacq et al. (2006) was cloned into a high-copy backbone (ColE1 origin of replication) carrying kanamycin resistance. The expression of the superfolder GFP reporter was placed under the control of the ptetO promoter. To control for plasmid copy number, RFP was constitutively expressed with the BBa_J23106 promoter and Plotkin RBS Kosuri et al. (2013) in a divergent orientation to sfGFP.

### Library synthesis

A comprehensive single-mutant TetR library was generated by replacing wildtype residues at positions 2–207 of TetR to all other 19 canonical amino acids (3,914 total mutant sequences). Oligonucleotides encoding each single-point mutation were synthesized as single-stranded Oligo Pools from Twist Bioscience and organized into 6 subpools, spanning six segments of the *tetRd* gene, corresponding to residues 2–39, 40–77, 78–115, 116–153, 154–191, and 192–207 of TetR(B) respectively. Additional sequence diversity was observed in the library due to error rates in the synthesis of single-stranded Oligo Pools, leading to the downstream identification of some double and triple mutant TetR variants. Oligo pools were encoded as a concatemer of the forward priming sequence, a BasI restriction site (5^′^-GGTCTC), six-base upstream constant region, TetR mutant sequence, six-base downstream constant region, a BsaI site (5^′^-GAGACC), and the reverse priming sequence. Subpools were resuspended in double-distilled water (ddH_2_ 0) to a final molal concentration of 25 ng/μL and amplified using primers specific to each oligonucleotide subpool with KAPA SYBR FAST qPCR (KAPA Biosystems; 1-ng template). A second PCR amplification was performed with KAPA HiFi (KAPA Biosystems; 1-μL qPCR template, 15 cycles maximum). We amplified corresponding regions of pSC101_TetR_specR with primers that linearized the backbone, added a BsaI restriction site, and removed the replaced wildtype sequence. Vector backbones were further digested with DpnI, BsaI, and Antarctic phosphatase before library assembly.

We assembled mutant libraries by combining the linearized sensor backbone with each oligo subpool at a molar ratio of 1:5 using Golden Gate Assembly Kit (New England Biolabs; 37 ^◦^C for 5 min and 60 ^◦^C for 5 min, repeated 30 times). Reactions were dialyzed with water on silica membranes (0.025-μm pores) for 1 h before transformed into DH10B cells (New England Biolabs). Library sizes of at least 100,000 colony-forming units (cfu) were considered successful. DH10B cells containing the reporter pColE1_sfGFP_RFP_kanR were transformed with extracted plasmids to obtain cultures of at least 100,000 cfu. Following co-transformation, the cultures for each subpool were stored as glycerol stocks and kept at -80 ^◦^C.

### Fluorescence-activated cell sorting

The subpool library cultures were seeded from glycerol stocks into 3 mL lysogeny broth (LB) containing 50 μg/mL kanamycin (kan) and 50 μg/mL spectinomycin (spec) and grown for 16 h at 37 ^◦^C. Each culture was then back-diluted into two wells of a 96-well plate containing LB kan/spec media using a dilution factor of 1:50 and grown for a period of 5 hours at 37 ^◦^C. Following incubation, each well containing saturated library culture was diluted 1:75 into 1× phosphate saline buffer (PBS), and fluorescence intensity was measured on a SH800S Cell Sorter (Sony). We first gated cells to remove debris and doublets and selected for variants constitutively expressing RFP. Using this filter, we then proceeded to draw an additional gate to select for mutants that displayed low fluorescence intensities in the absence of anhydrotetracycline (aTC). This gate allowed us to select for mutants which retained the ability to repress GFP expression in the presence of an inducer, using prior measurements of WT TetR(B) as a reference for selecting repression competent mutants Leander et al. (2020). Utilizing these gates, we sorted 500,000 events for each gated population and recovered these cells in 5 mL of LB at 37 ^◦^C before adding 50 μg/mL of kan and spec. Following the addition of antibiotics, the cultures were incubated at 37 ^◦^C for 16 hours.

Following overnight growth, each culture was back diluted into two wells of a 96-well plate containing LB kan/spec media using a 1:50 dilution factor. Upon reaching an OD600 ∼ 0.2, aTC was added to one of the two wells at a final concentration of 1 μM, with the well not receiving aTC serving as the uninduced control population. Cells were then incubated for another 4-hour growth period at 37 ^◦^C before being diluted into 1x PBS using a dilution factor of 1:75, and fluorescence intensity was measured on a SH800S Cell Sorter.

We first gated cells to remove debris and doublets and selected for variants constitutively expressing RFP before obtaining the distribution of fluorescence intensities (FITC-A) across the uninduced and induced populations of each of the 6 subpools (***Figure 3***—figure Supplement 8). Using the uninduced control population of each subpool as a reference, gates were drawn to capture mutants which displayed no response to the presence of aTC in the induced populations. Using these gates, 500,000 events were sorted from the induced populations of each subpool and cells were subsequently recovered at 37 ^◦^C using 5 mL of LB media before being plated onto LB agar plates containing 50 μg/mL kan and spec.

### Clonal screening of dead mutants

To screen for functionally deficient variants of TetR, 100 individual colonies were picked from each subpool and grown to saturation in a 96-well plate for 6 hours. Saturated cultures were then diluted 1:50 in LB-kan/spec media and grown in the presence and absence of 1 μM aTC for 6 hours before OD600 and GFP fluorescence (Gain: 40; Excitation: 488/20; Emission: 525/20) were read on a multiplate reader (Synergy HTX, BioTek). Fluorescence was normalized to OD600, and the fold inductions of each clone were calculated by dividing its normalized fluorescence in the presence of inducer by the normalized fluorescence in the absence of inducer. The fold-induction of WT TetR was tested in parallel with these screens to serve as a benchmark for function, and clones which displayed < 50% activity to WT were replated and validated in triplicate before being sent for sanger sequencing (Functional Biosciences).

### aTC dose response measurements

TetR mutants identified during clonal screening were reinoculated into 3 mL of LB Kan/Spec media and grown overnight. After an overnight growth period, each culture was added to 4 rows of 12 wells in a 96-well plate containing LB Kan/Spec media using a dilution factor of 1:50. Each mutant was tested across 12 different concentrations of aTC in quadruplicate fashion, with each row having an identical concentration gradient across the 12 wells. The aTC concentration gradient across the 12 wells were 0 nM, 10 nM, 20 nM, 40 nM, 60 nM, 80 nM, 100 nM, 150 nM, 250 nM, 500 nM, 750 nM, and 1μM aTC.

Following reinoculation into 96-well plates, selected mutants were placed on a plate shaker set to 900 RPM and incubated at 37 ^◦^C for 5 hours. Following this 5-hour growth period, the plates were removed from the plate shaker and the OD600 and GFP fluorescence (Gain: 40; Excitation: 488/20; Emission: 525/20) of each well was read using a multiplate reader (Synergy HTX, BioTek). Fluorescence was normalized to OD600 for each well. The normalized fluorescence at each concentration was measured for 4 replicates of each mutant (unless otherwise stated). The fluorescence of GFP reporter-only control (no TetR), grown under identical conditions, was measured in the same way. This reporter-only control served as an upper bound to the fluorescence that could be measured in the experiment and was accompanied by a WT TetR-GFP reporter control, used to certify the precision of measurements made at different times.

### Synthesis of combined TetR mutants

To assess the predictive capability of the additive model, eight combined mutants were selected to undergo aTC dose response characterization. These combined mutants were synthesized using clonal DNA fragments from Twist Biosciences coding for the amino acid sequence of each of the selected combined mutants. The gene fragments were individually cloned into sensor plasmid backbones identical to those used in the previous identification of non-functional TetR(B) mutants following the manufacturers protocol for Gibson Assembly (New England Biolabs). The newly constructed plasmids carrying each of the combined mutants were then transformed into E. coli (DH10B) cells containing a GFP reporter plasmid identical to that used in the previous characterization of non-functional TetR(B) mutants. After a 1-hour recovery period, 100 μL of recovered cells were plated onto LB kan/spec agar plates for each of the combination mutants and incubated for 16 hours at 37 ^◦^C.

Following overnight growth, individual colonies were picked and inoculated into 3 mL of LB kan/spec media to be grown overnight at 37 ^◦^C. Following overnight growth, samples from each culture were submitted for sanger sequencing to confirm the identity of each of the synthesized combined mutants. Following sequence verification, the aTC dose-response behavior of each combined mutant was characterized using an experimental setup consistent with dose-response measurements made for the previously identified TetR(B) mutants.

### Model parameter estimation

As described above, the estimation of the three biophysical parameters (ɛ_D_, ɛ_L_ and γ) for each mutant is divided into two steps. Firstly, as ɛ_L_ and γ do not affect F C at *c* = 0 (***Equation 1***), we directly calculate the value of ɛ_D_ from the leakiness of an induction curve, which is measured with high precision in our experiments. With the calculated ɛ_D_ (ɛ_D_ ^w*T*^ is set to 0 as reference), we then fit the full induction curve to obtain the values of ɛ_L_ and γ using the method of Bayesian inference. Here, we first construct the statistical model that describes the data generating process based on ***Equation 1***, and specify the prior distributions of the relevant parameters. This enables the derivation of the conditional probability of the parameter values of a mutant given its induction data, known as the posterior distribution. The posterior distribution is then sampled using Markov chain Monte Carlo (MCMC), from which we infer the values of ɛ_L_ and γ directly. The validity of the statistical model and computational algorithm is fully tested with several metrics, and all details regarding the parameter estimation process briefed here are provided in the Supplementary file.

## Code and data availability

All code and data used in this work can be accessed at the Github repository https://github.com/liuzhbu/Two_Domain_Allostery.

## Author contributions

Z.L., S.R. and Q.C. conceptualized the work. Z.L. and T.G. conducted the research. Z.L. and T.G. analyzed the data. All authors are involved in writing the manuscript.

## Acknowledgments

This work is funded by NIH Director’s New Innovator Award DP2GM132682 (SR) and Shaw Scientist Award (SR), and R35-GM141930 (QC). Development of the Bayesian inference model was partially supported by grant ML-21-016 from the Dreyfus foundation (QC). Computational resources from the Extreme Science and Engineering Discovery Environment (XSEDE), which is supported by NSF grant number ACI-1548562, are greatly appreciated; part of the computational work was performed on the Shared Computing Cluster which is administered by Boston University’s Research Computing Services (URL: https://www.bu.edu/tech/ support/research/).

---

**Figure 2—figure supplement 1.**
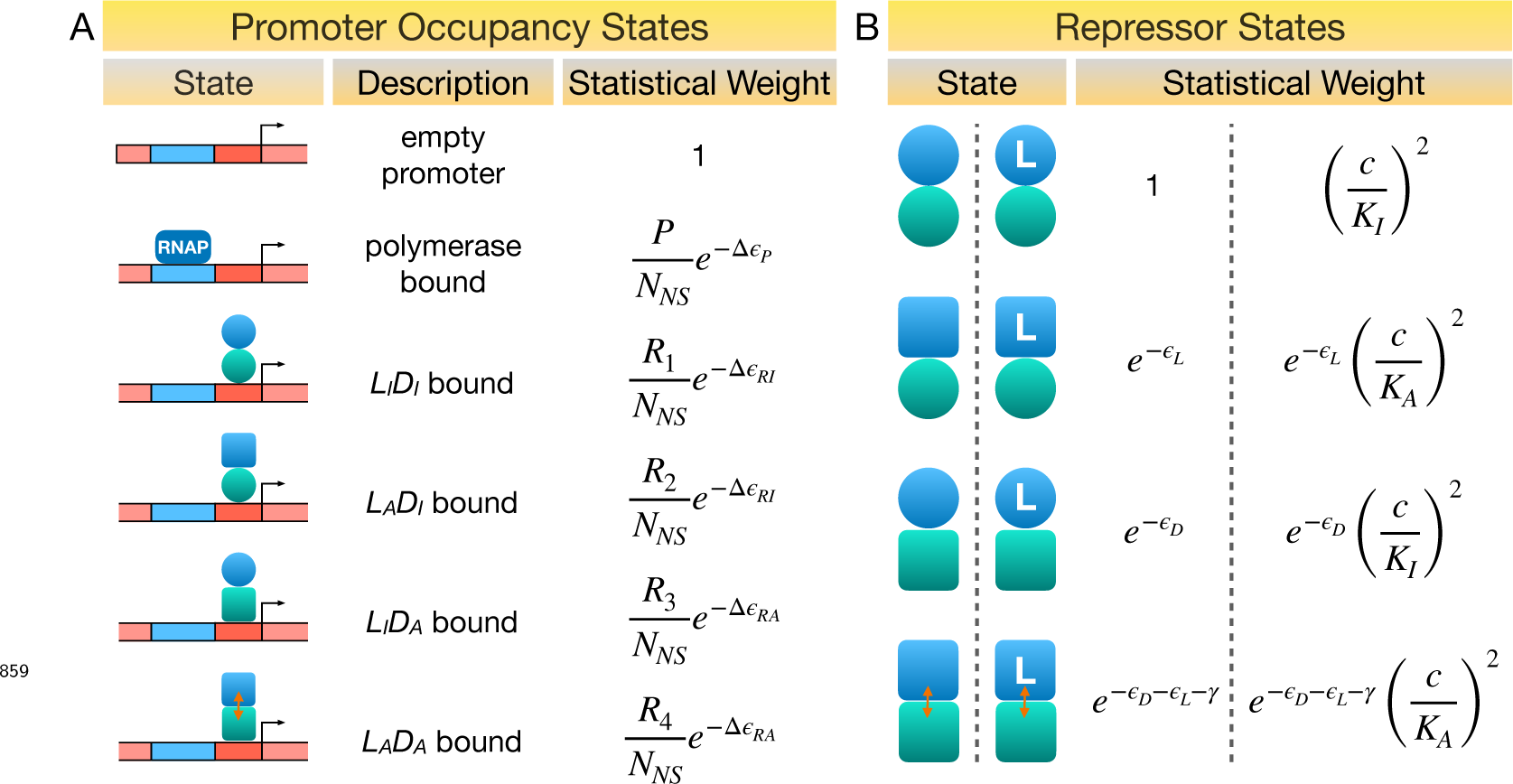
Statistical weights of promoter occupancy states and repressor states. (A) Statistical weights of the promoter occupancy states, with the empty promoter state taken as reference. *P* is the average number of RNAP per cell. *R*_1_, *R*_2_, *R*_3_ and *R*_4_ denote the average number of repressors in the L_I_ D_I_, L_A_ D_I_, L_I_ D_A_ and L_A_ D_A_ state per cell, respectively. *N*_*N*s_ is the number of non-specific DNA binding sites in the cell. Δɛ_*P*_, Δɛ_*R*A_ and Δɛ_*R*I_ represent the energy differences between specific and non-specific DNA binding of RNAP, repressor with DBD in the active and inactive conformations, respectively. (B) Statistical weights of the allosteric states of the repressor, with the L_I_ D_I_ state taken as reference. *K*_A_ and *K*_I_ are the dissociation constants of ligand to the repressor with LBD in the active and inactive conformations, respectively, and *c* is ligand concentration. Partial binding of ligand is ignored in the symmetric model. All energy terms in the exponents are evaluated in the unit of *k*_B_ *T*.

**Figure 2—figure supplement 2.**
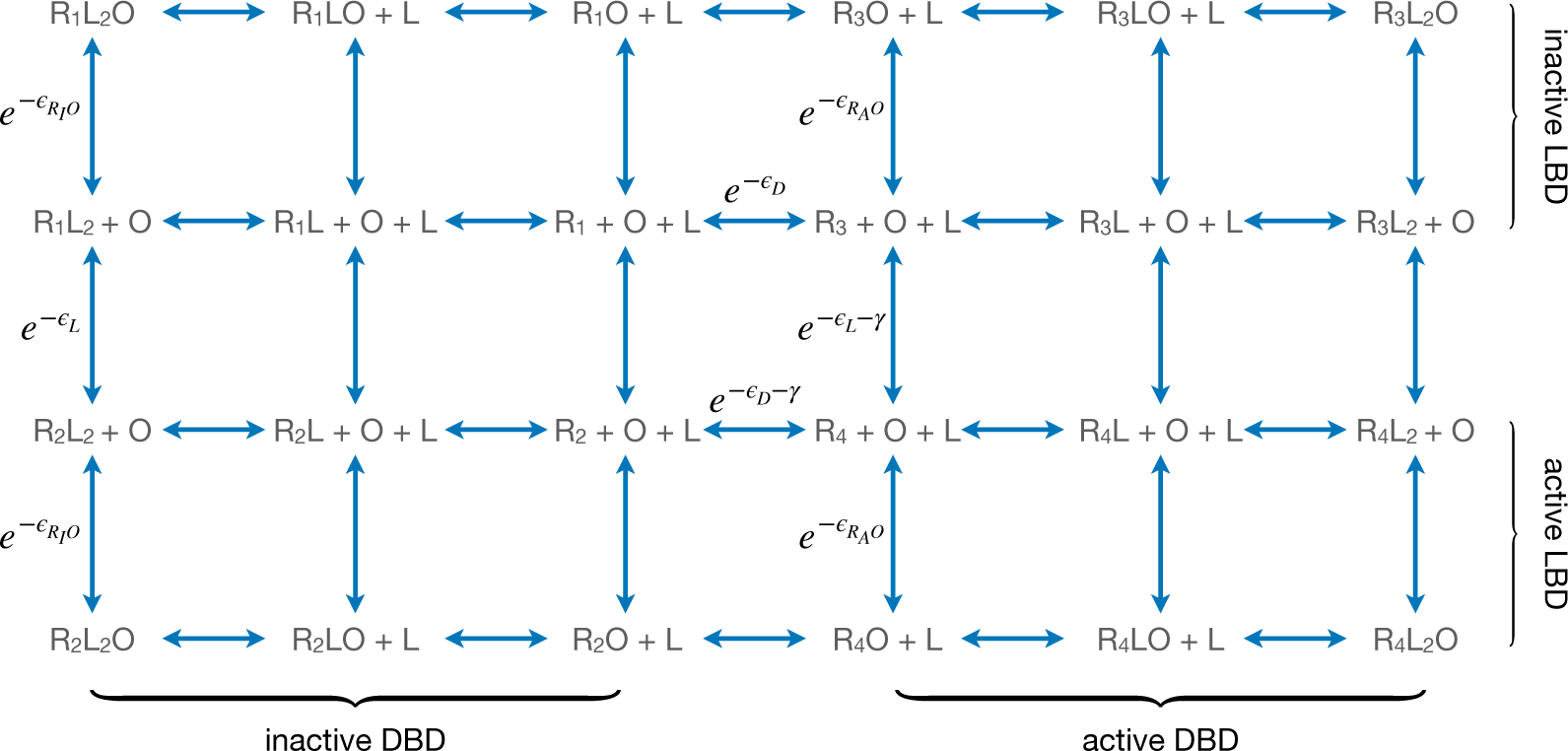
Equilibria among different conformational and binding states of the repressor. Here, L and O represent ligand and operator respectively. ɛ_*R*0_ and ɛ_*R*0_ are the A I free energies of operator binding for repressor with DBD in the active and inactive conformation, respectively. *R*_1_, *R*_2_, *R*_3_ and *R*_4_ denote the L_I_ D_I_, L_A_ D_I_, L_I_ D_A_ and L_A_ D_A_ state of the repressor, respectively. All energy terms in the exponents are evaluated in the unit of *k*_B_ *T*.

**Figure 2—figure supplement 3.**
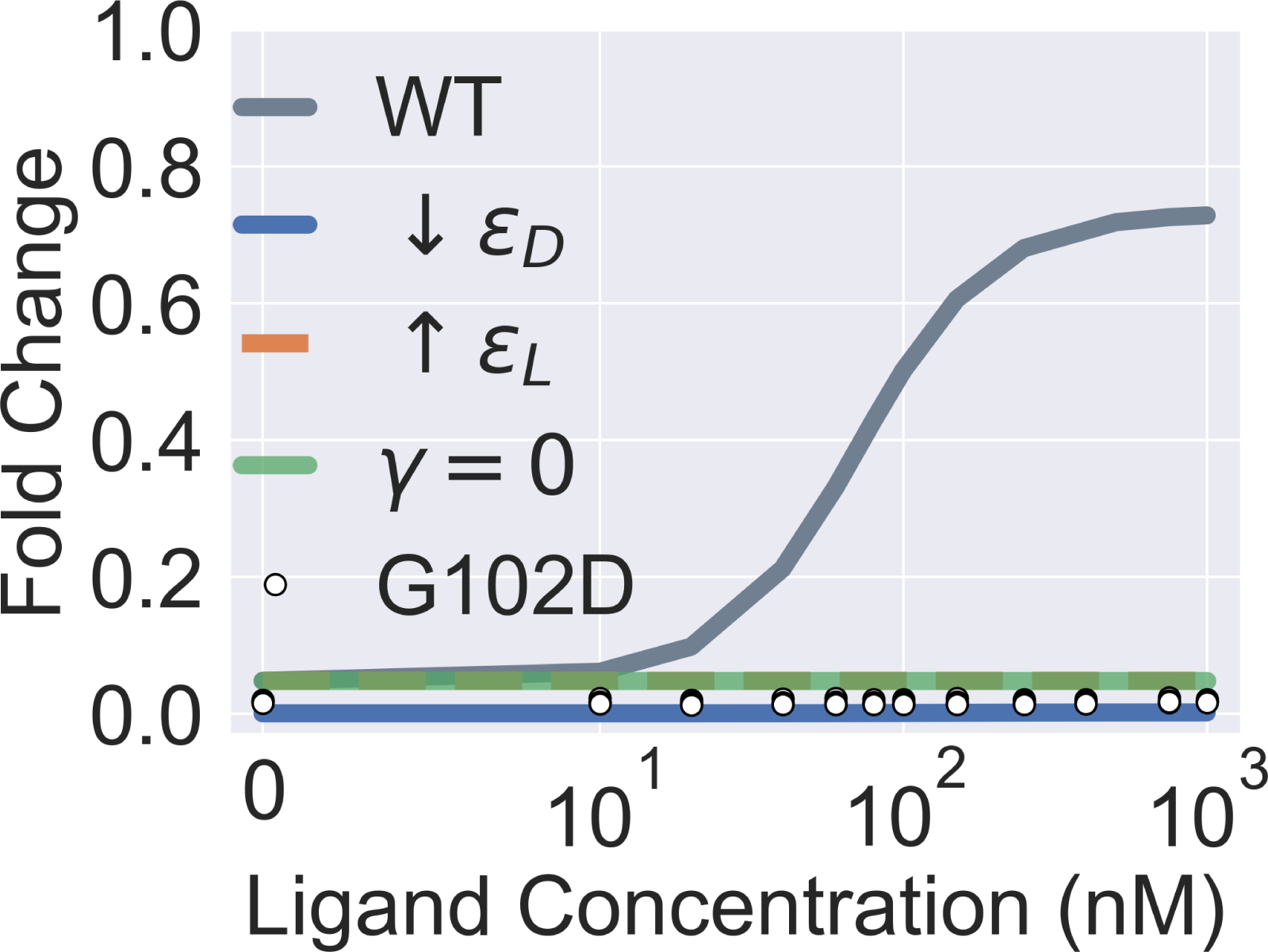
Extended parametric study of main text ***Equation 1***. The four colored curves show that a flat induction curve can result from tuning each one of the three main biophysical parameters of the two-domain model alone (by decreasing ɛ_D_ or increasing ɛ_L_ from the WT value, or by setting γ to 0). The white points show the experimental induction data for the mutant G102D (measurements of 4 biological replicates at each ligand concentration). Note that the colored curves in the figure are generated for better visualization of the model parameters’ effect on the induction curve and not to be compared with the experimental data of G102D. The leakiness of G102D (0.0173) is higher than that of the WT (0.0086) determined in experiments.

**Figure 3—figure supplement 1.**
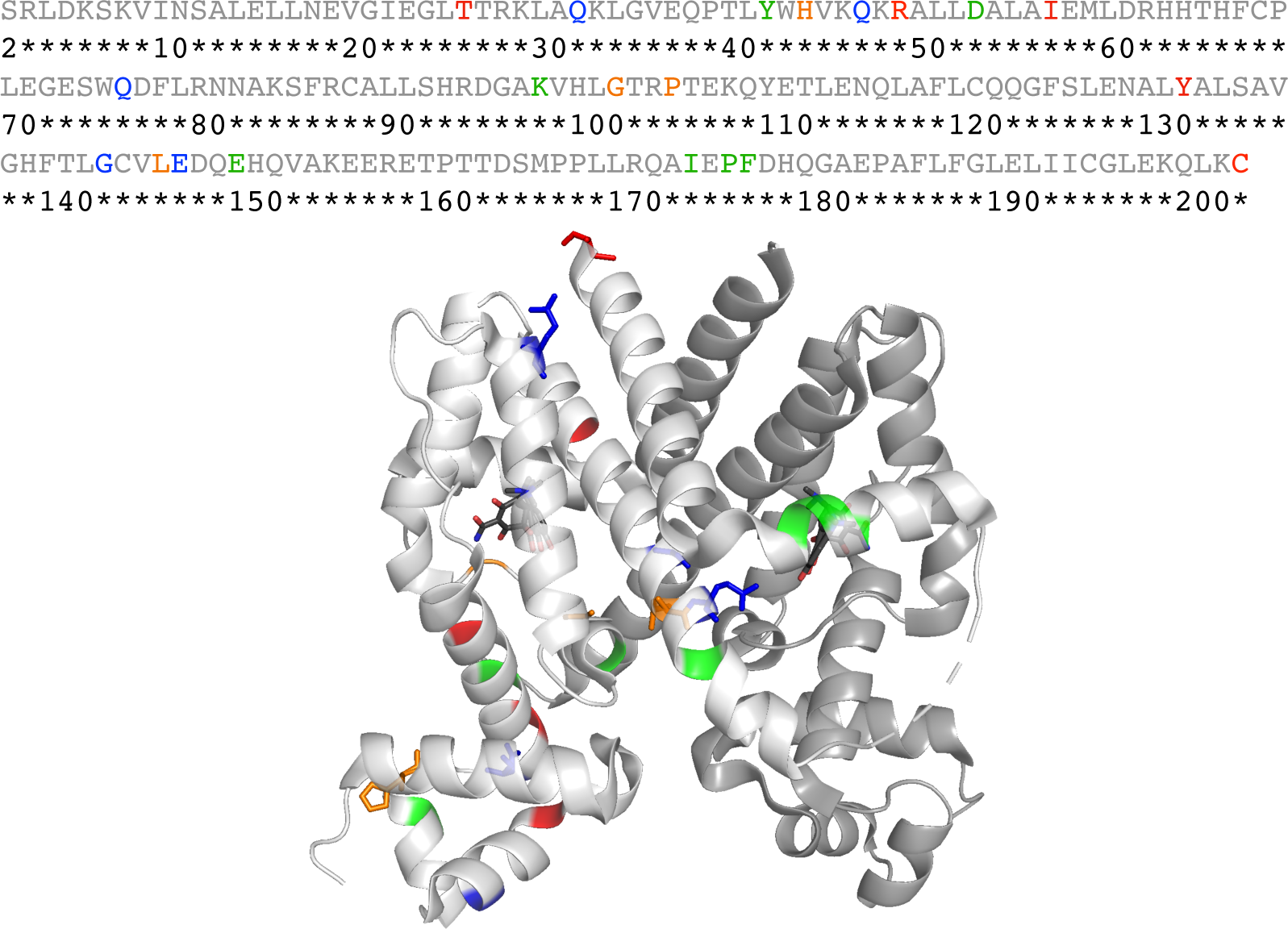
Sequence and structural distributions of the 21 residues chosen for the mutation analyses in this work. The upper panel shows the sequence of TetR (residue 2-203), where the 21 residues chosen for mutation analyses in this work are colored red, orange, green or blue (while the other residues are colored grey). The lower panel shows the crystal structure of TetR(B) in complex with minocycline and magnesium (PDB code: 4AC0). Here, the two identical monomers of TetR are colored white and grey respectively, while the residues chosen for mutation analyses are colored in the same way as in the sequence above in the white monomer. Specifically, the 5 red residues from top to bottom are C203, Y132, I57, R49 and T26. The 4 orange residues from left to right are H44, P105, G102 and L146. The 5 blue residues from top to bottom are Q76, G143, E147, Q47 and Q32. The 7 green residues from left to right are Y42, D53, K98, E150, F177, P176 and I174. Some of the 21 residues are presented in the stick format to aid visualization while all other residues are presented in the cartoon format.

**Figure 3—figure supplement 2.**
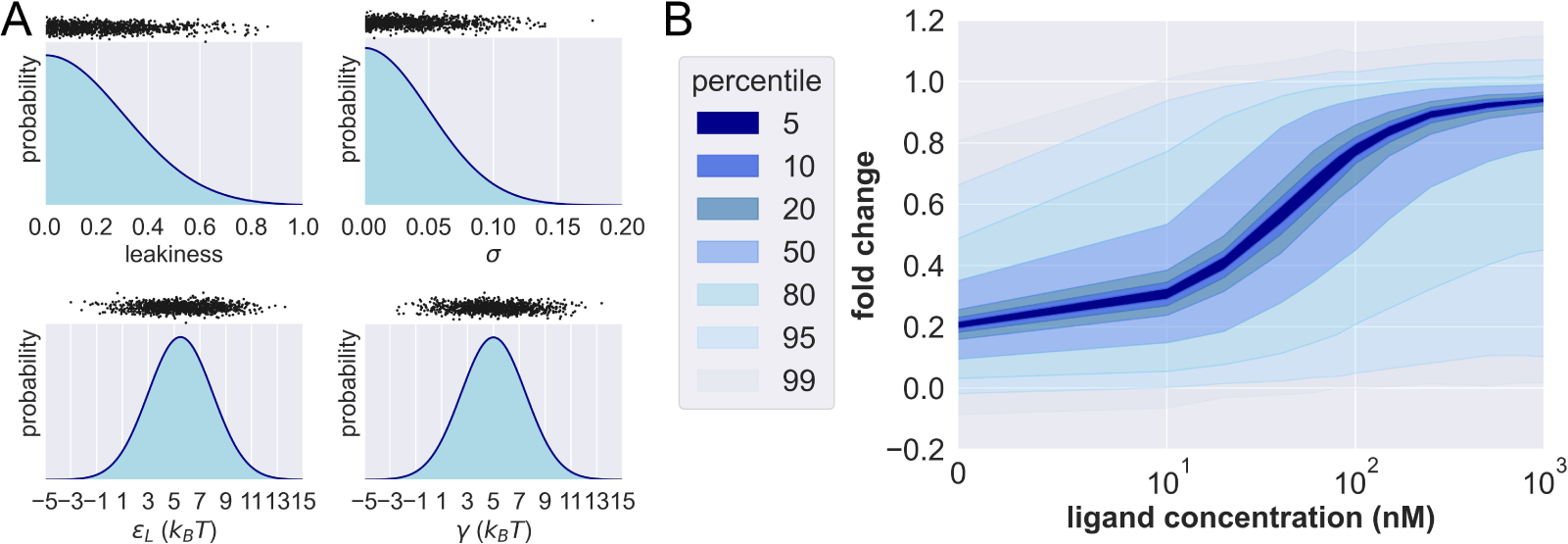
Prior probability distributions and prior predictive check. (A) Density functions of the prior distributions of leakiness, σ, ɛ_L_ and γ. The black dots above the prior distributions show the 1000 prior predictive draws of the corresponding parameter. (B) Percentiles of the simulated fold changes using the 1000 sets of parameters shown in (A).

**Figure 3—figure supplement 3.**
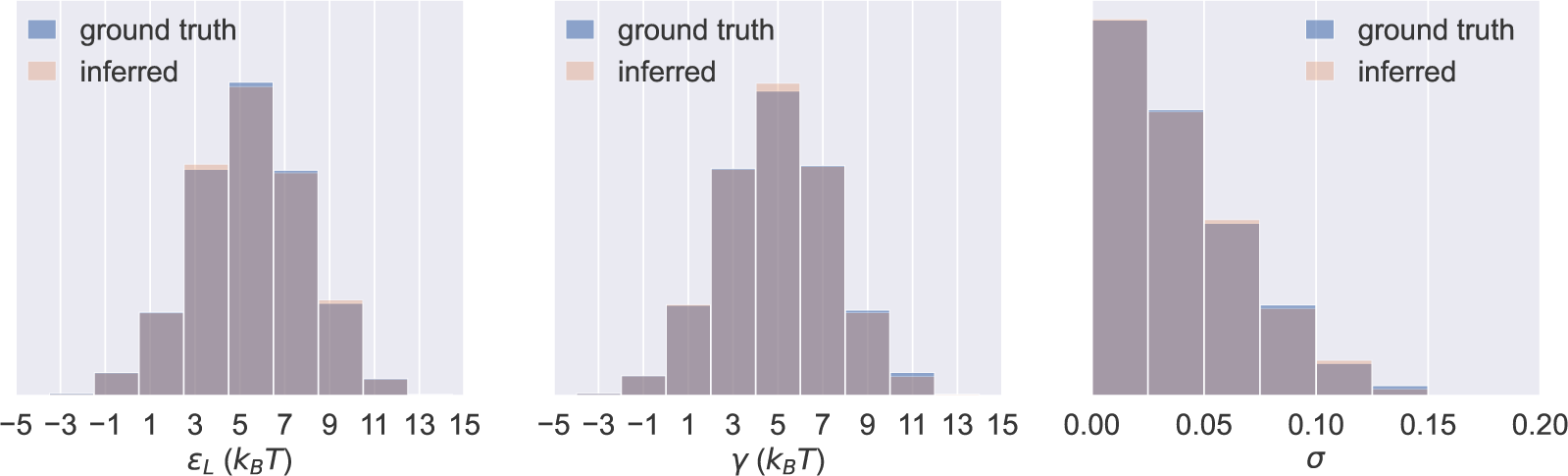
Probability distributions of ɛ_L_, γ and σ values in the 1000 sets of prior predictive draws (ground truth) and the average of the corresponding 1000 sets of inferred posterior distributions of the parameters (inferred).

**Figure 3—figure supplement 4.**
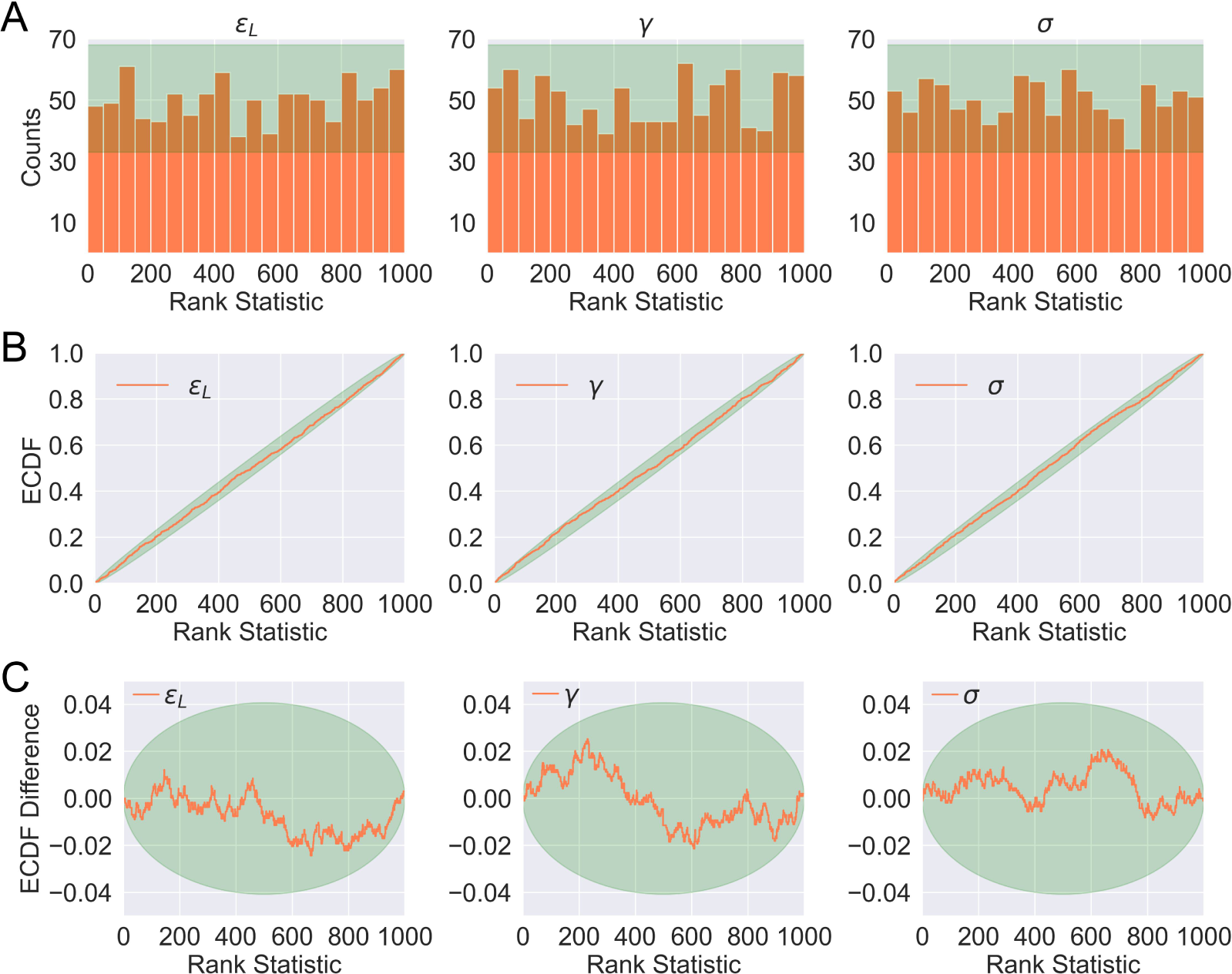
Distributions of rank statistics of the prior predictive draws relative to the corresponding posterior samples. (A) Histograms (20 bins); (B) ECDF plots; (C) ECDF difference plots of rank statistics of ɛ_L_, γ and σ. The green bands in (A)-(C) show the 99^*th*^percentile expected from a true uniform distribution.

**Figure 3—figure supplement 5.**
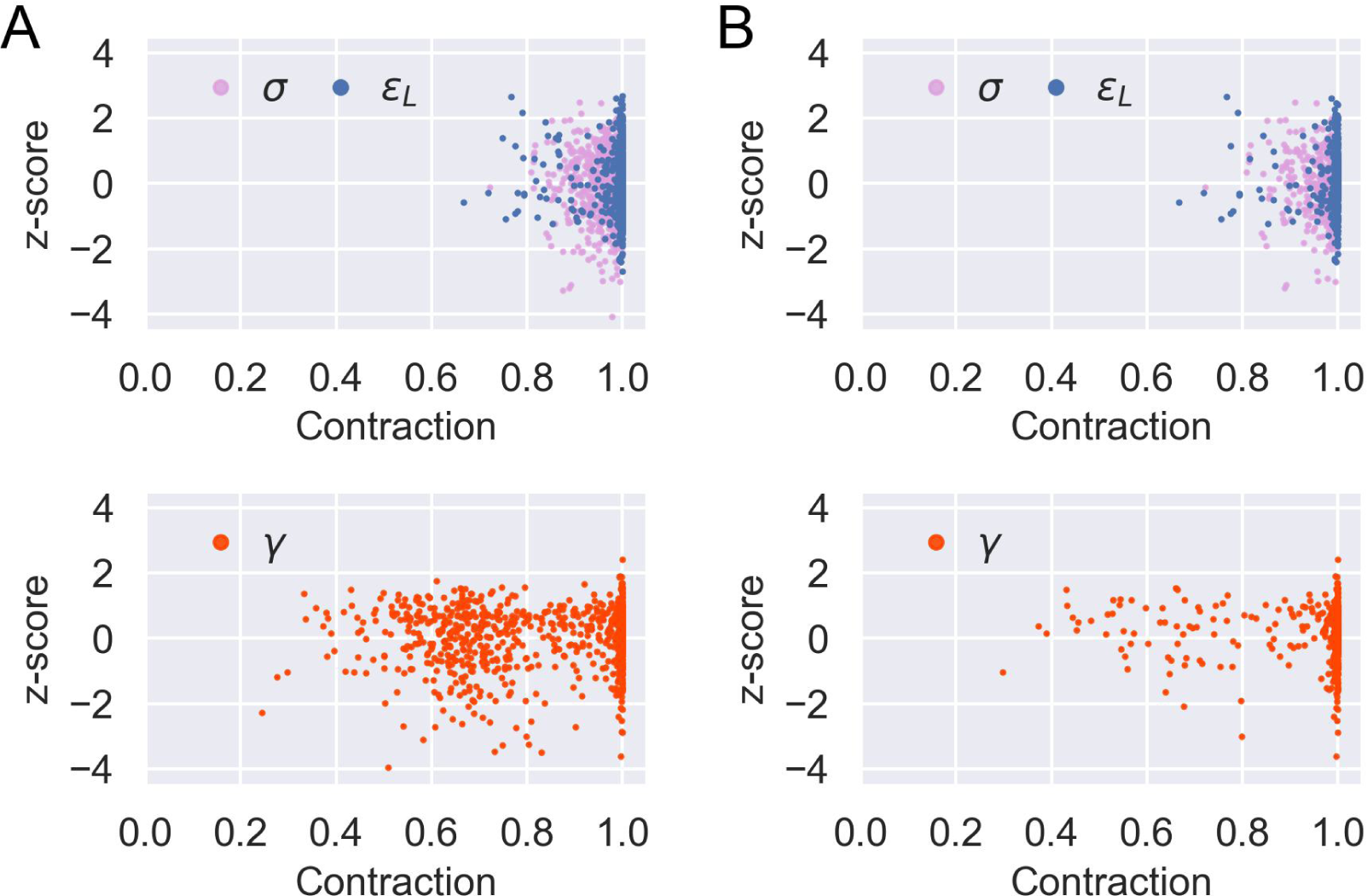
Sensitivity analysis for model parameter inference. Posterior z-score and posterior contraction of the inferred posterior distribution for each of the 1000 prior predictive draws of parameters and the corresponding simulated data except: (A) parameters of flat induction curves (see Supplementary file section 3 and Eq. (32)); (B) parameters of flat induction curves or μ(*c*=1000 nM)>0.97 (see Supplementary file section 3 and Eq. (34)). Accordingly, there are 970 and 523 data points for each parameter in (A) and (B), respectively.

**Figure 3—figure supplement 6.**
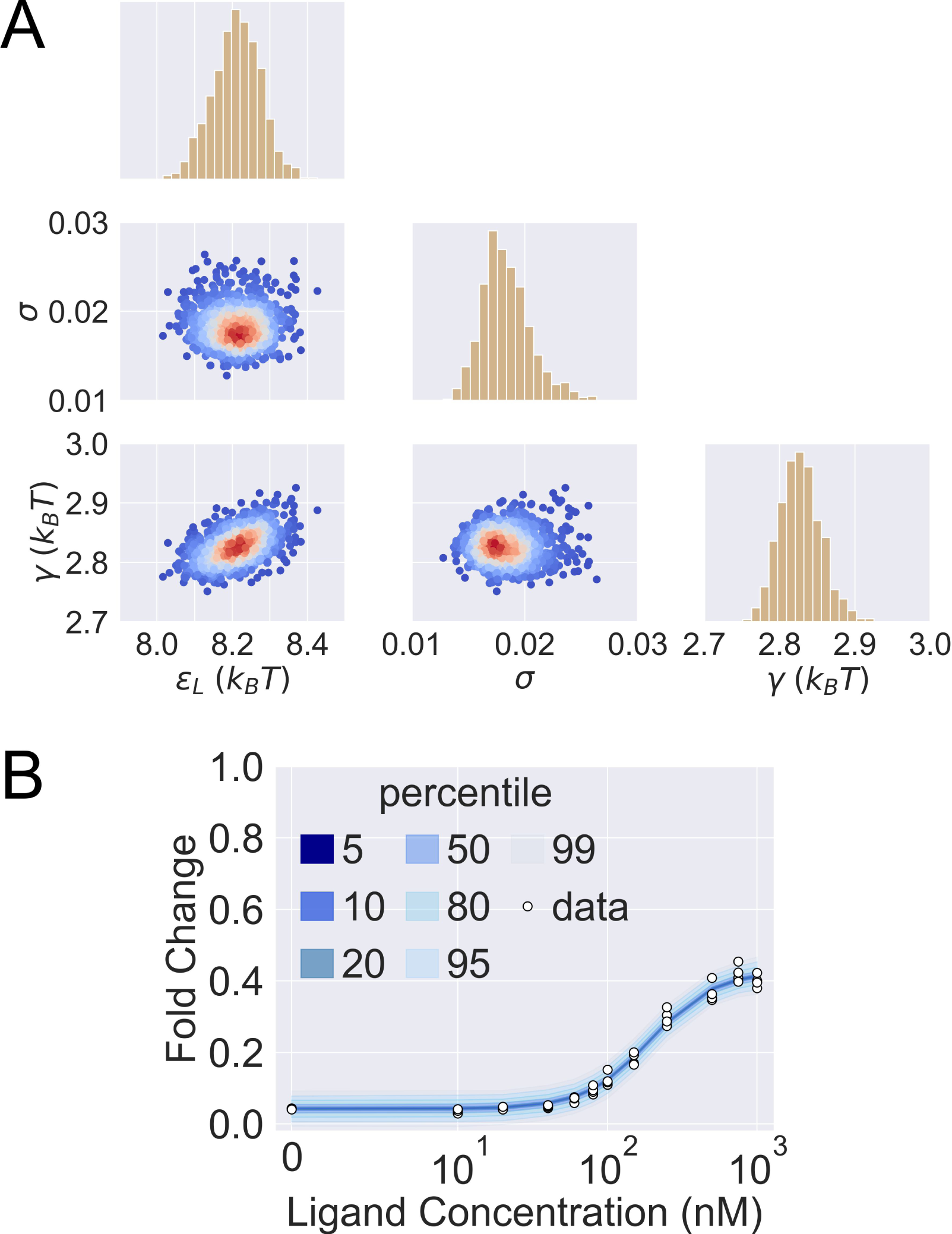
Posterior predictive check of mutant G102D-Y42M-I57N. (A) 1000 sets of posterior samples of ^{^ɛ, γ, σ^}^. The scattered plots show the joint distributions of the parameters, colored by the log probability of the parameter combinations. Blue/red corresponds to low/high probability. The histograms show the marginal distributions of the corresponding individual parameters. (B) Percentiles of the simulated fold change measurements using the 1000 sets of posterior samples based on Supplementary file Eq. (24). The white data points show the experimental induction data of four biological replicates.

**Figure 3—figure supplement 7.**
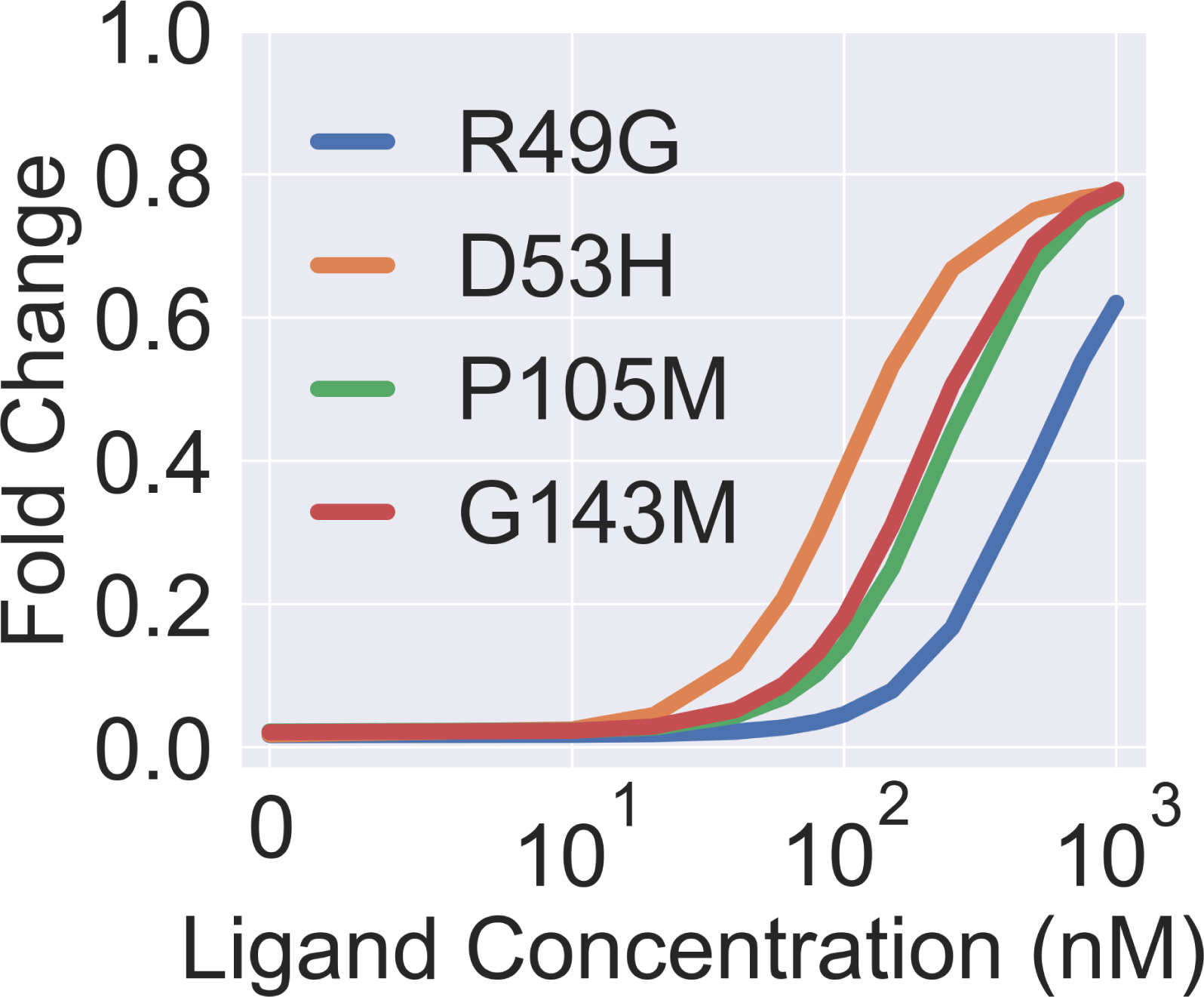
Theoretical induction curves of the four dead mutants when their γ values are set to the WT value while using their respective ɛ_D_ and ɛ_L_ values (taken from Figure 3B in the main text). The theoretical induction curve is calculated with main text ***Equation 1***

**Figure 3—figure supplement 8.**
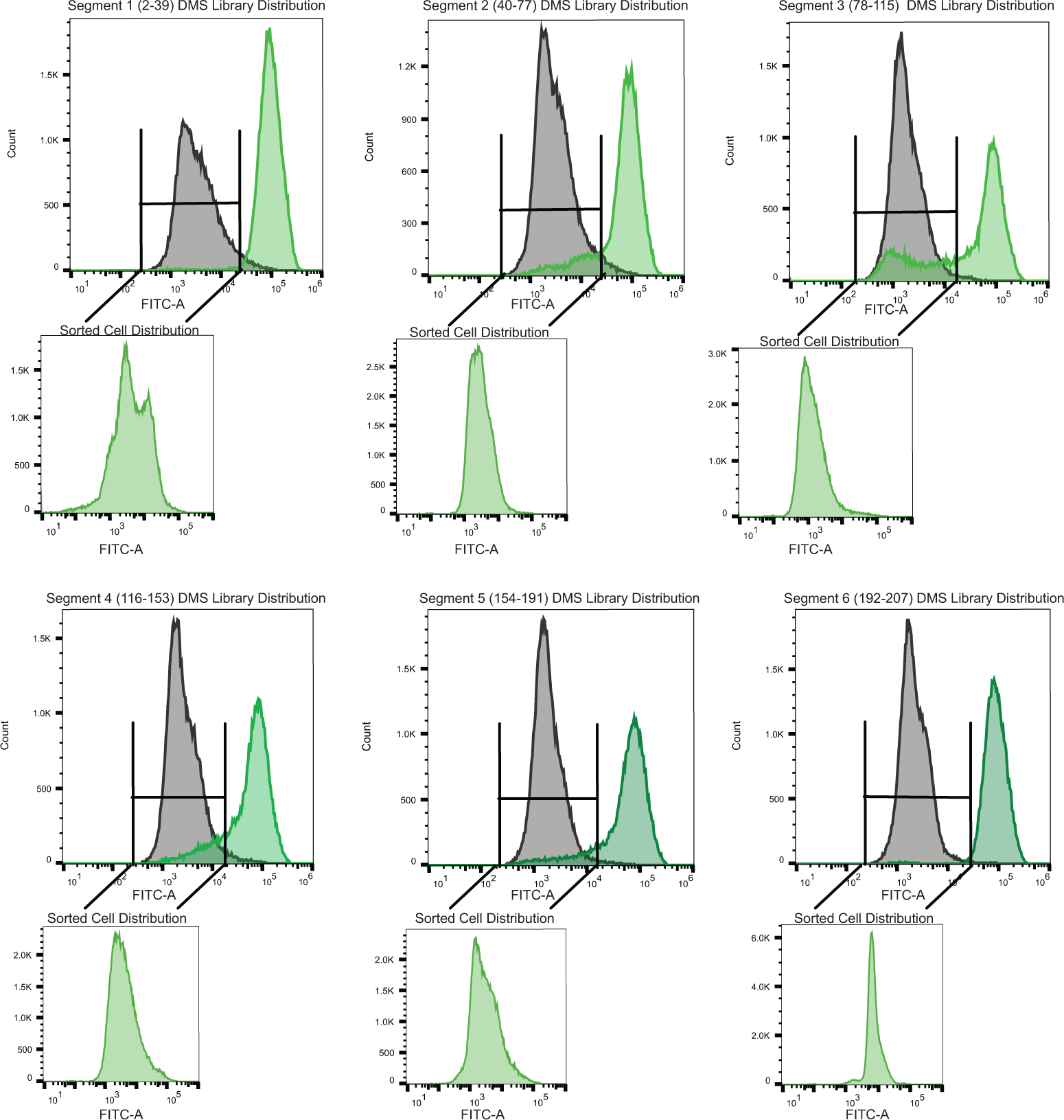
Sorting scheme to identify dead variants. Each panel is a tiled segment of single site saturation mutants of defined length along the TetR gene. The residue number spanning each segment is shown above the panel. The gray distribution denotes the uninduced population of cells containing mutants. The dark green distribution on the same panel denotes the population of cells responding to the anhydrotetracycline. The uninduced population of cells containing mutants when was added was sorted and reflowed which is shown outside the main panel as the light green distribution.

**Figure 4—figure supplement 1.**
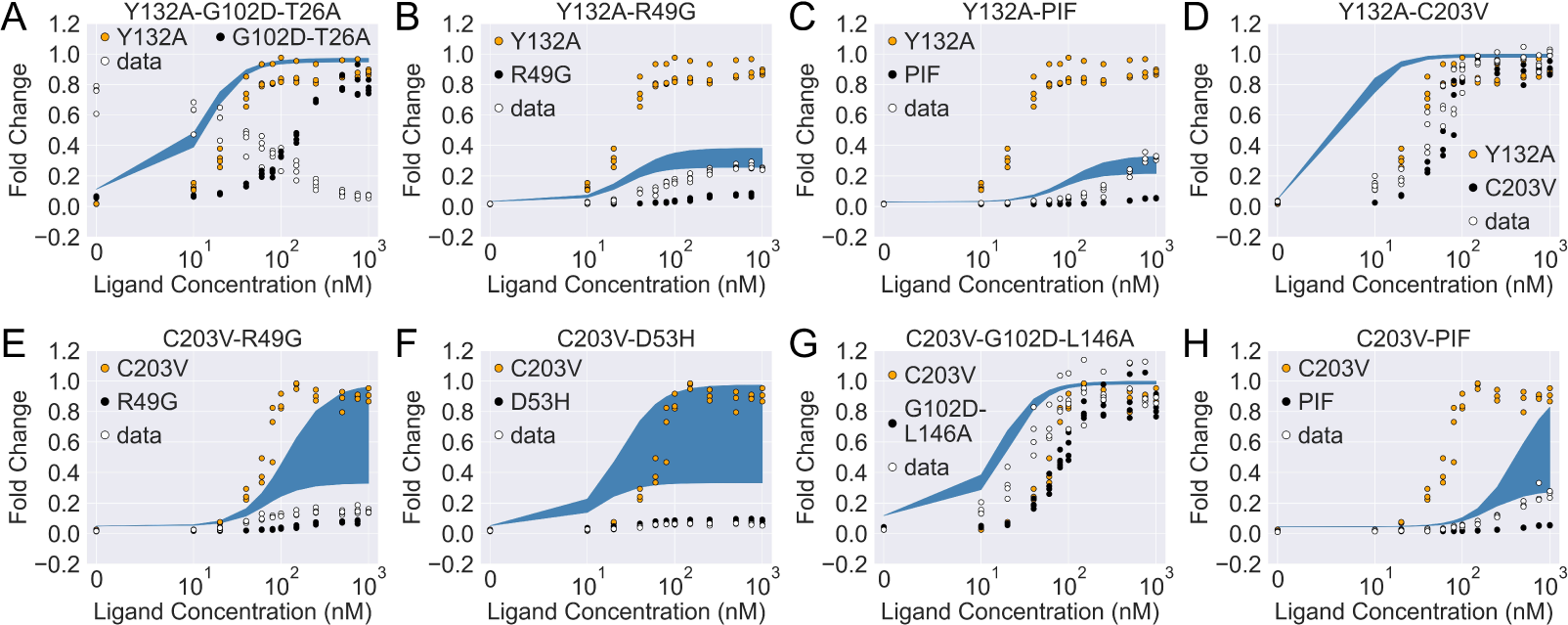
The induction curves of the eight combined mutants calculated using the basic additive model (i.e., with α_1,*p*_ = α_2,*p*_ = 1 in main text ***Equation 4***). In each plot, the white data points show the experimental induction data of the combined mutants, while the black and orange ones show data of the individual mutants. The blue band show the 95^*th*^percentile of the induction curve prediction from the additive model.

**Figure 4—figure supplement 2.**
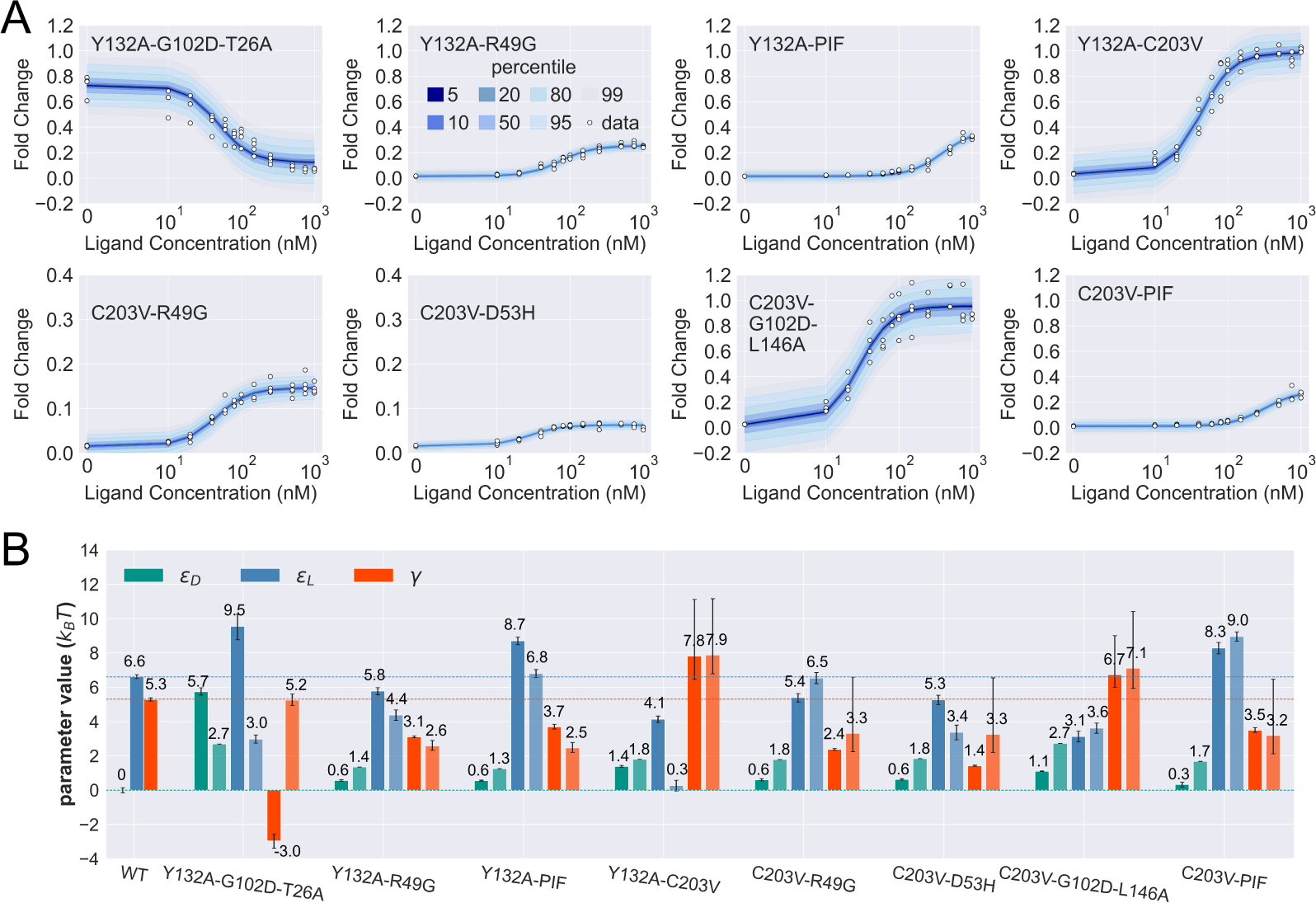
Induction curves of the eight combined mutants and the corresponding parameter estimation results as well as the basic additive model predictions. (A) Percentiles of the simulated fold change measurements using the inferred posterior parameters of each mutant based on Supplementary file Eq. (24). The white data points show the experimental induction data of four biological replicates (three replicates for C203V-PIF). (B) The inferred parameter values of WT and the 8 combined mutants. Error bars of ɛ_L_ and γ represent the upper and lower bounds of the 95 percent credible region. For each combined mutant, the parameters shown by the lighter and darker bar plots are the results from the basic additive model and direct fitting, respectively. The horizontal lines show the WT parameter values for reference.

