## Supplementary file for "A parametrized two-domain thermodynamic model explains diverse mutational effects on protein allostery"

**Zhuang Liu**[1], **Thomas Gillis**[2], **Srivatsan Raman**[2,3,4], **Qiang Cui**[1,5*]

[1]Department of Physics, Boston University, Boston, United States; [2]Department of Biochemistry, University of Wisconsin, Madison, United States; [3]Department of Chemistry, University of Wisconsin, Madison, United States; [4]Department of Bacteriology, University of Wisconsin, Madison, United States; [5]Department of Chemistry, Boston University, Boston, United States

### 1. Derivation of main text Equation 1

#### For nonzero ligand concentration

Eq. **1** of main text expresses the gene expression level rescaled by that of an unregulated promoter (fold change) as a function of system and allosteric parameters, whose detailed derivation is given below. Following Griffin et al. ***Chure et al.*** (***2019***) and previous works on thermodynamic models of transcription ***Ackers et al.*** (***1982***); ***Buchler et al.*** (***2003***); ***Vilar and Leibler*** (***2003***); ***Bintu et al.*** (***2004***); ***Kuhlman et al.*** (***2007***); ***Daber et al.*** (***2011***), the gene expression level is considered to be proportional to the probability of the promoter being bound by RNA polymerase (RNAP). Accordingly, the possible occupancy states of the promoter and their corresponding statistical weights are listed in main text **Figure 2—figure Supplement 1**A, from which we can calculate the probability of the RNAP bound state as its statistical weight divided by the partition function as given in Eq. (1).

$$p_{RNAP}^{bound} = \frac{\frac{P}{N_{NS}} e^{-\Delta\epsilon_P}}{1 + \frac{P}{N_{NS}} e^{-\Delta\epsilon_P} + \frac{R_1}{N_{NS}} e^{-\Delta\epsilon_{RI}} + \frac{R_2}{N_{NS}} e^{-\Delta\epsilon_{RI}} + \frac{R_3}{N_{NS}} e^{-\Delta\epsilon_{RA}} + \frac{R_4}{N_{NS}} e^{-\Delta\epsilon_{RA}}} \tag{1}$$

Here, $P$ is the average number of RNAP per cell. $R_1$, $R_2$, $R_3$ and $R_4$ are the average numbers of repressors in the $L_I D_I$, $L_A D_I$, $L_I D_A$ and $L_A D_A$ state per cell, respectively. $N_{NS}$ is the number of non-specific DNA binding sites in the cell, which is taken to be the length of the Escherichia Coli (E. Coli) genome in base pairs ($\sim 4.6\times10^6$) ***Chure et al.*** (***2019***). All RNAP and repressors are assumed to be bound to either the specific or non-specific binding sites on DNA ***Chure et al.*** (***2019***); ***Garcia and Phillips*** (***2011***); ***Rünzi and Matzura*** (***1976***); ***von Hippel et al.*** (***1974***); ***Kao-Huang et al.*** (***1977***); ***Normanno et al.*** (***2015***); ***Stracy et al.*** (***2021***); ***von Hippel and Berg*** (***1989***), where the specific binding sites of repressor and RNAP are the operator and promoter sequence, respectively (such assumption is not essential for the derivation of main text **Equation 1** ***Vilar and Leibler*** (***2003***), see the last part of this section). When a repressor binds to the operator (orange rectangle in main text **Figure 2—figure Supplement 1**A), it obstructs RNAP from binding to the adjacent promoter (blue rectangle in main text **Figure 2—figure Supplement 1**A) and the expression of downstream genes (right pink rectangle in main text **Figure 2—figure Supplement 1**A). $\Delta\epsilon_P$, $\Delta\epsilon_{RA}$ and $\Delta\epsilon_{RI}$ represent the energy differences between specific and non-specific DNA binding of RNAP, repressor with

DBD in the active and inactive conformations, respectively. Note that the free energy quantities are all measured in the unit of $k_B T$, thus the Boltzmann factors throughout our discussion do not include $k_B T$ explicitly.

The statistical weight of a given promoter occupancy state is evaluated based on its energy and the number of microscopic states it corresponds to. With the empty promoter taken as the reference state (statistical weight equals 1), we can then calculate the statistical weights of the other operator states as their probabilities relative to the reference state. Since $N_{NS} \gg R$ ($R = \sum_{i=1}^{4} R_i$ $\sim$ 5000 is the average number of all repressors in the cell) and $P$ ($P \approx 1000$) ***Klumpp and Hwa*** (***2008***); ***Kosuri et al.*** (***2013***), the non-specific binding of repressors and RNAP are considered to be independent. When the promoter is occupied by a RNAP, there are $\binom{N_{NS}}{P-1}$ ways to arrange the remaining $(P-1)$ RNAPs on the $N_{NS}$ nonspecific binding sites. Thus, the probability of RNAP bound state relative to the empty promoter state (all $P$ RNAPs are bound to nonspecific binding sites) is given by:

$$\frac{p_{RNAP}^{bound}}{p^{empty}} = \frac{\binom{N_{NS}}{P-1}}{\binom{N_{NS}}{P}} e^{-\Delta\varepsilon_P}, \tag{2}$$

where the energy difference between the two promoter occupancy states is accounted for in the Boltzmann weight. As $N_{NS} \gg P$, the right-hand side (RHS) of Eq. (2) can be easily simplified to $\frac{P}{N_{NS}} e^{-\Delta\varepsilon_P}$, which is the statistical weight of the RNAP-bound promoter state given in Eq. (1) and main text **Figure 2—figure Supplement 1**A. The statistical weights of the other repressor-bound states of promoter are calculated following the same reasoning. It's noted that, since the number of promoter-operator sequences present in a cell ($\sim$ 10-20) is much smaller than $P$ and $R$ (see the main text section "$Materials\ and\ methods$"), the binding status of each promoter is regarded to be independent of others ***Brewster et al.*** (***2014***).

Next, the $p_{RNAP}^{bound}$ of a repressor-regulated promoter is divided by that of an unregulated promoter (where $R$=0) to get the relative gene expression level, which is defined as fold change ($FC$) in main text **Equation 1**.

$$FC = \frac{p_{RNAP}^{bound}(R>0)}{p_{RNAP}^{bound}(R=0)} = \frac{1 + \frac{P}{N_{NS}} e^{-\Delta\varepsilon_P}}{1 + \frac{P}{N_{NS}} e^{-\Delta\varepsilon_P} + \frac{R_1+R_2}{N_{NS}} e^{-\Delta\varepsilon_{RI}} + \frac{R_3+R_4}{N_{NS}} e^{-\Delta\varepsilon_{RA}}} \tag{3}$$

With the assumption of weak promoter $1 \gg \frac{P}{N_{NS}} e^{-\Delta\varepsilon_P}$, and that the energy difference between specific and non-specific DNA binding of repressor with DBD in the inactive conformation ($\Delta\varepsilon_{RI}$) is small ***Chure et al.*** (***2019***), Eq. (3) can be simplified to

$$FC = \frac{1}{1 + \frac{R_3+R_4}{N_{NS}} e^{-\Delta\varepsilon_{RA}}} = \frac{1}{1 + \frac{R}{N_{NS}} \left\{ p_{R_3}(c) + p_{R_4}(c) \right\} e^{-\Delta\varepsilon_{RA}}}. \tag{4}$$

Here, $p_{R_3}(c)$ and $p_{R_4}(c)$ are the probabilities of a repressor being in the $L_I D_A$ and $L_A D_A$ state at a given ligand concentration $c$, respectively, which can be calculated from the statistical weights of the possible repressor states. As shown in main text **Figure 2—figure Supplement 1**B, the unbound $L_I D_I$ state is taken as the reference state with a statistical weight of 1. $\varepsilon_L$ ($\varepsilon_D$) is the free energy increase of the repressor when LBD (DBD) transition from the inactive to the active conformation, and $\gamma$ is the additional free energy penalty when both domains adopt the active conformations. $K_A$ and $K_I$ are the dissociation constants of ligand to the repressor with LBD in the active and inactive conformations, respectively.

Similar to the calculation of $p_{RNAP}^{bound}$, the sum of $p_{R_3}(c)$ and $p_{R_4}(c)$ is then given by

$$p_{R_3}(c) + p_{R_4}(c) = \frac{e^{-\varepsilon_D} \left\{ 1 + \left(\frac{c}{K_I}\right)^2 \right\} + e^{-\varepsilon_D - \varepsilon_L - \gamma} \left\{ 1 + \left(\frac{c}{K_A}\right)^2 \right\}}{1 + \left(\frac{c}{K_I}\right)^2 + e^{-\varepsilon_L} \left\{ 1 + \left(\frac{c}{K_A}\right)^2 \right\} + e^{-\varepsilon_D} \left\{ 1 + \left(\frac{c}{K_I}\right)^2 \right\} + e^{-\varepsilon_D - \varepsilon_L - \gamma} \left\{ 1 + \left(\frac{c}{K_A}\right)^2 \right\}}. \tag{5}$$

For the simplicity and mechanistic clarity of the model, we assume that ligand binding to the repressor with LBD in the inactive conformation is negligible ($c/K_I \ll 1$ for $c$ below 1 $\mu$M, the maximum ligand concentration used in our experiments). Then plugging Eq. (5) back into Eq. (4) we have

$$FC = \frac{1}{1 + \frac{Re^{-\Delta\varepsilon_{RA}}}{N_{NS}} e^{-\varepsilon_D} \frac{1+e^{-\varepsilon_L-\gamma}\left\{1+\left(\frac{c}{K_A}\right)^2\right\}}{1+e^{-\varepsilon_L}\left\{1+\left(\frac{c}{K_A}\right)^2\right\}+e^{-\varepsilon_D}+e^{-\varepsilon_D-\varepsilon_L-\gamma}\left\{1+\left(\frac{c}{K_A}\right)^2\right\}}}. \tag{6}$$

We can state that $c/K_A \gg 1$ for all ligand concentrations where we measured gene expression levels experimentally (10 nM$\leq c \leq$1000 nM) besides $c$=0, based on the apparent dissociation constants of the ligand (anhydrotetracycline; aTC) binding to TetR reported in the literature ***Scholz et al.*** (***2000***); ***Schubert et al.*** (***2004***) and the assumption that $e^{-\varepsilon_L} \ll 1$. Thus, at the nonzero ligand concentrations where we measured gene expression levels experimentally, Eq. (6) can be simplified to

$$FC = \frac{1}{1 + \frac{Re^{-\Delta\varepsilon_{RA}}}{N_{NS}} e^{-\varepsilon_D} \frac{1+e^{-\varepsilon_L-\gamma}\left(\frac{c}{K_A}\right)^2}{1+e^{-\varepsilon_L}\left(\frac{c}{K_A}\right)^2+e^{-\varepsilon_D}+e^{-\varepsilon_D-\varepsilon_L-\gamma}\left(\frac{c}{K_A}\right)^2}}. \tag{7}$$

Further assuming $e^{-\varepsilon_D} \ll 1$ and $e^{-\varepsilon_D-\gamma} \ll 1$, Eq. (7) is reduced to

$$FC = \frac{1}{1 + \frac{Re^{-\Delta\varepsilon_{RA}}}{N_{NS}} e^{-\varepsilon_D} \frac{1+e^{-\varepsilon_L-\gamma}\left(\frac{c}{K_A}\right)^2}{1+e^{-\varepsilon_L}\left(\frac{c}{K_A}\right)^2}}, \tag{8}$$

which is the same as the main text **Equation 1** after substituting $\frac{Re^{-\Delta\varepsilon_{RA}}}{N_{NS}}$ and $K_A$ with $R^*$ and $K$, respectively, to simplify notations.

All energy terms in the exponents are evaluated in the unit of $k_BT$ in this work, and we've assumed that $e^{-\varepsilon_D}$, $e^{-\varepsilon_D-\gamma}$, $e^{-\varepsilon_L}$ and $e^{-\varepsilon_L-\gamma}$ to be much smaller than one based on the consideration that free energy differences between different conformations of a TetR like protein is usually on the order of a few $k_BT$ ***Chure et al.*** (***2019***); ***Motlagh et al.*** (***2014***).

### For zero ligand concentration

At c=0, Eq. (6) becomes

$$FC = \frac{1}{1 + \frac{Re^{-\Delta\varepsilon_{RA}}}{N_{NS}} e^{-\varepsilon_D} \frac{1+e^{-\varepsilon_L-\gamma}}{1+e^{-\varepsilon_L}+e^{-\varepsilon_D}+e^{-\varepsilon_D-\varepsilon_L-\gamma}}}. \tag{9}$$

Under the same set of assumptions stated above, Eq. (9) is reduced to $FC = (1 + R^* e^{-\varepsilon_D})^{-1}$, which agrees with Eq. (8) at $c$=0. It should also be noted that $\varepsilon_D$ plays a dominant role in deciding $FC$ at $c$=0 (leakiness of the induction curve) compared with $\varepsilon_L$ and $\gamma$, even based on the full functional form of Eq. (9). Thus, the difference between the leakiness of the induction curves of different mutants analyzed in this study is attributed solely to their difference in $\varepsilon_D$, as mutations of operator-binding residues are avoided and $R^*$ is taken to be constant across these mutants (see the section "*Model parameter estimation*" below).

### Inclusion of single-ligand-bound state of repressor

As shown in the main text section "*Overview of the two-domain thermodynamic model of allostery*" and main text **Figure 2—figure Supplement 1**, single-ligand-bound state of the repressor is ignored in our symmetric two-domain model and the derivation of Eq. (8) (main text **Equation 1**) for simplicity. Nonetheless, including single-ligand-bound repressor states in the derivation above leads to the same form of Eq. (8), which we demonstrate below. Allowing single ligand binding, Eq. (5) becomes

$$p_{R_3}(c) + p_{R_4}(c) = \frac{e^{-\varepsilon_D}\left(1+\frac{c}{K_I}\right)^2 + e^{-\varepsilon_D-\varepsilon_L-\gamma}\left(1+\frac{c}{K_A}\right)^2}{\left(1+\frac{c}{K_I}\right)^2 + e^{-\varepsilon_L}\left(1+\frac{c}{K_A}\right)^2 + e^{-\varepsilon_D}\left(1+\frac{c}{K_I}\right)^2 + e^{-\varepsilon_D-\varepsilon_L-\gamma}\left(1+\frac{c}{K_A}\right)^2}, \quad (10)$$

while Eq. (1)-(4) stay unchanged. Accordingly, Eq. (6) becomes

$$FC = \frac{1}{1 + \frac{Re^{-\Delta\varepsilon_{RA}}}{N_{NS}} e^{-\varepsilon_D} \frac{1+e^{-\varepsilon_L-\gamma}\left(1+\frac{c}{K_A}\right)^2}{1+e^{-\varepsilon_L}\left(1+\frac{c}{K_A}\right)^2+e^{-\varepsilon_D}+e^{-\varepsilon_D-\varepsilon_L-\gamma}\left(1+\frac{c}{K_A}\right)^2}}, \quad (11)$$

which is reduced to the same form of Eq. (6) and Eq. (9) at $c/K_A \gg 1$ and $c = 0$, respectively. The subsequent steps in the derivation that lead to Eq. (8) then stay the same as before.

### Alternative derivation of main text equation 1

Eq. **1** of main text can be derived from a different perspective of the system as well, which is based on the equilibria among different conformational and binding states of the repressor as specified in main text **Figure 2—figure Supplement 2** ***Daber et al.*** (***2011***). As gene expression level is considered proportional to the probability that a promoter is bound by RNAP, it can be alternatively written as

$$FC = \frac{p_{RNAP}^{bound}(R > 0)}{p_{RNAP}^{bound}(R = 0)} = \frac{[O]}{[O] + [R_1O] + [R_2O] + [R_3O] + [R_4O]}. \quad (12)$$

Here, [O] is the concentration of free operator in the cell, and $[R_1O]$ to $[R_4O]$ are the concentrations of operator bound repressor that is in the corresponding conformations (main text **Figure 2—figure Supplement 2**) regardless of ligand binding states (e.g., $[R_1O]$ is the total concentration of species $[R_1O]$, $[R_1LO]$ and $[R_1L_2O]$). Based on the equilibrium constants given in main text **Figure 2—figure Supplement 2**, we then have

$$FC = \frac{1}{1 + ([R_1] + [R_2])e^{-\varepsilon_{R_IO}} + ([R_3] + [R_4])e^{-\varepsilon_{R_AO}}}, \quad (13)$$

where $[R_1]$ to $[R_4]$ are the concentrations of the repressor in the corresponding conformations divided by 1M (not bound to operator) regardless of its ligand binding states. Under the condition that the total number of repressors is much larger than that of operators (or promoters) in the cell ($[R_{tot}] \gg [O_{tot}]$), Eq. (13) can be rewritten in a more familiar form as given below.

$$FC = \frac{1}{1 + [R_{tot}]e^{-\varepsilon_{R_AO}}\{f + s(1-f)\}} \quad (14)$$

$$f = \frac{e^{-\varepsilon_D}\left(1+\frac{c}{K_I}\right)^2 + e^{-\varepsilon_D-\varepsilon_L-\gamma}\left(1+\frac{c}{K_A}\right)^2}{\left(1+\frac{c}{K_I}\right)^2 + e^{-\varepsilon_L}\left(1+\frac{c}{K_A}\right)^2 + e^{-\varepsilon_D}\left(1+\frac{c}{K_I}\right)^2 + e^{-\varepsilon_D-\varepsilon_L-\gamma}\left(1+\frac{c}{K_A}\right)^2} \quad (15)$$

$$s = \frac{e^{-\varepsilon_{R_IO}}}{e^{-\varepsilon_{R_AO}}} \quad (16)$$

Assuming that the operator binding affinity of a repressor with active DBD is much higher than that of a repressor with inactive DBD ($s \ll 1$), such that the effect of inactive operator binding on gene expression is negligible, Eq. (14) can be further simplified to

$$FC = \frac{1}{1 + [R_{tot}]e^{-\varepsilon_{R_AO}} \frac{e^{-\varepsilon_D}\left(1+\frac{c}{K_I}\right)^2+e^{-\varepsilon_D-\varepsilon_L-\gamma}\left(1+\frac{c}{K_A}\right)^2}{\left(1+\frac{c}{K_I}\right)^2+e^{-\varepsilon_L}\left(1+\frac{c}{K_A}\right)^2+e^{-\varepsilon_D}\left(1+\frac{c}{K_I}\right)^2+e^{-\varepsilon_D-\varepsilon_L-\gamma}\left(1+\frac{c}{K_A}\right)^2}}. \quad (17)$$

With the same assumption used in the derivation from Eq. (5) to (6) in the last section ($c/K_I \ll 1$ for $c$ below 1 $\mu$M, the maximum ligand concentration used in our experiments), we have

$$FC = \frac{1}{1+[R_{tot}]e^{-\epsilon_{RAO}}e^{-\epsilon_D}\frac{1+e^{-\epsilon_L-\gamma}\left(1+\frac{c}{K_A}\right)^2}{1+e^{-\epsilon_L}\left(1+\frac{c}{K_A}\right)^2+e^{-\epsilon_D}+e^{-\epsilon_D-\epsilon_L-\gamma}\left(1+\frac{c}{K_A}\right)^2}}. \tag{18}$$

Eq. (18) is the same as Eq. (11) only with $\frac{Re^{-\Delta\epsilon_{RA}}}{N_{NS}}$ ($R^*$ of main text **Equation 1**) redefined as $[R_{tot}]e^{-\epsilon_{RAO}}$, and thus we'll arrive at the main text **Equation 1** following the same derivation steps leading from Eq. (11) to (8).

Note that the difference in the interpretation of the $R^*$ term in main text **Equation 1** obtained from the derivations adopting the two perspectives demonstrated above doesn't affect our inference results based on the two-domain allosteric model (see the section "*Model parameter estimation*" below).

### 2. Extended parametric study of main text Equation 1

#### Effect of $\epsilon_L$ on induction curve

As demonstrated in the main text section "*System-level ramifications of the two-domain model*" and main text **Figure 2***A*, increasing $\epsilon_L$ from the wildtype (WT) value alone can result in an induction curve featuring a sharply varying tail instead of the full sigmoidal shape (purple curve of main text **Figure 2***A*). Further increasing $\epsilon_L$ however, will result in a flat induction curve (orange curve of main text **Figure 2—figure Supplement 3**), as the free energy gain of ligand binding becomes too limited to cause any noticeable effect on the allosteric response of the repressor even at its highest concentration used in our experiments ($c = 1000$ nM). Theoretically, based on main text **Equation 1**, an induction curve should always maintain the sigmoidal shape however high $\epsilon_L$ becomes, given that we extend the concentration range for induction data measurement so that main text **Equation 2** is still applicable. Yet in practice, we find Escherichia coli (E. coli) stops growing normally at $c$(aTC)>1000 nM, which is thus set as our concentration limit for induction data measurement.

#### Effect of $\epsilon_D$ on induction curve and the parametric degeneracy of flat induction curve

As demonstrated in the main text section "*System-level ramifications of the two-domain model*" and main text **Figure 2***B*, decreasing $\epsilon_D$ from the WT value alone simultaneously lowers the leakiness and the level of saturation of the induction curve while maintaining the sigmoidal shape (purple curve of main text **Figure 2***B*). However, further decrease of $\epsilon_D$ in principle can reduce their difference (dynamic range) to a level below the detection limit of our experiments as shown in Eq. (19), and the induction curve will appear essentially flat (blue curve of main text **Figure 2—figure Supplement 3**).

$$\lim_{c\to\infty} FC - FC(c=0) = \frac{R^*(1-e^{-\gamma})}{(1+R^*e^{-\epsilon_D-\gamma})(1+R^*e^{-\epsilon_D})}e^{-\epsilon_D} \tag{19}$$

A flat induction curve can also result from vanishing inter-domain coupling ($\gamma$=0) where $FC$ becomes a constant $1/(1+R^*e^{-\epsilon_D})$ independent of $c$ (main text **Equation 1**), as shown by the green curve of main text **Figure 2—figure Supplement 3**. Therefore, we've demonstrated the degeneracy of flat induction curves in the parameter space of the two-domain model, which prevent an accurate characterization of mutants with such induction curves (e.g., G102D, see main text **Figure 2—figure Supplement 3**).

#### $\gamma$ and the monotonicity of $FC(c)$

From main text **Equation 1**, we have

$$\frac{dFC(c)}{dc} = 2\frac{FC^2}{\left(1+e^{-\epsilon_L}\frac{c^2}{K^2}\right)^2}\frac{R^*e^{-\epsilon_D-\epsilon_L}c}{K^2}(1-e^{-\gamma}). \tag{20}$$

As the fraction term of the RHS of Eq. (20) is positive besides at $c = 0$ (where it equals 0), the derivative of $FC$ with respect to $c$ is $\geq 0$, $\leq 0$ and =0 when $\gamma$>0, $\gamma$<0 and $\gamma$=0, respectively for all ligand concentrations.

### 3. Model parameter estimation

In this work, the estimation of allosteric parameters ($\epsilon_D$, $\epsilon_L$ and $\gamma$) of a mutant is divided into two steps: 1. Calculation of $\epsilon_D$ from the leakiness of its induction curve, which is then taken as a constant in step two; 2. Evaluation of $\epsilon_L$ and $\gamma$ based on the full induction curve using the method of Bayesian inference. Detailed procedures are provided below.

#### Calculation of $\epsilon_D$

As shown in main text **Equation 1**, the leakiness of an induction curve is determined by $R^*$ and $\epsilon_D$ of the corresponding mutant, where $R^*$, specified by Eq. (8) and (18), is determined by system constants (e.g., $N_{NS}$ or cell volume), repressor copy number per cell (taken to be a constant across mutants ***Chure et al.*** (***2019***)), and the affinity between operator and the repressor with an active DBD. Therefore, since no mutation of direct DNA-interacting residues is made in the 24 mutants investigated in this work ***Ramos et al.*** (***2005***), the variation of induction curve leakiness of a mutant repressor from the WT is attributed solely to the change of $\epsilon_D$. This enables the direct calculation of $\epsilon_D$ of the mutants from their induction curve leakiness as given in Eq. (21).

$$\epsilon_D = -ln\left(\frac{1}{FC(c=0)} - 1\right) + ln(R^*) \tag{21}$$

Although the exact value of $R^*$ is not resolved in our experiments, it doesn't affect the evaluation of the change of $\epsilon_D$ in a mutant relative to the WT (main text **Figure 3** and **Figure 4—figure Supplement 2**). This reflects how the mutations modify the allosteric nature of the repressor, which is what we focus on here (also see the next part of this section).

In main text **Figure 3** and **Figure 4—figure Supplement 2**, the reported value of $\epsilon_D$ is calculated based on the average of $FC(c = 0)$ of four or more biological replicates for each mutant (except G102D-HQQ, C203V and C203V-PIF, which have three replicates), with the reference $\epsilon_D$(WT) set to 0. Error bar of $\epsilon_D$ is calculated based on the standard error of the mean (SEM) of the corresponding leakiness measurement. The uncertainty of $\epsilon_D$ is well below 0.1 $k_BT$ for all mutants except for C203V-PIF and Y132A-G102D-T26A, whose $\epsilon_D$ uncertainties are 0.15 $k_BT$ and 0.2 $k_BT$, respectively. Thus, based on the characteristic effects of different allosteric parameters on the induction curve as established in main text **Equation 1**, the evaluation of $\epsilon_D$ is decoupled from that of $\epsilon_L$ and $\gamma$. This helps resolving the phenotype degeneracy in the parameter space of the two-domain model, enabling their estimation with high precision.

#### Estimation of $\epsilon_L$ and $\gamma$ with Bayesian inference

$\epsilon_L$ and $\gamma$ of mutants are estimated simultaneously based on the induction curves following the standard workflow of Bayesian inference (see below) as introduced in previous works ***Chure et al.*** (***2019***); ***Schad et al.*** (***2020***). The distinct effects of $\epsilon_L$ and $\gamma$ on the induction curve (see main text section "*System-level ramifications of the two-domain model*") ensure their evaluation with a low level of uncertainty.

##### i. Building a generative statistical model

In Bayesian inference, we want to estimate the value of $\epsilon_L$ and $\gamma$ of a mutant given its induction data. To do this, we first need a statistical model that describes the conditional probability of different parameter values given the experimental observation (induction curve in this work). According to Bayes' theorem, such conditional probability (known as posterior distribution) is calculated as

$$p(\epsilon_L, \gamma | y) = \frac{f(y|\epsilon_L, \gamma)g(\epsilon_L)g(\gamma)}{\iint d\epsilon_L d\gamma\ f(y|\epsilon_L, \gamma)g(\epsilon_L)g(\gamma)}. \tag{22}$$

Here, $y$ is the experimental data (fold change); $f(y|\varepsilon_L, \gamma)$ calculates the likelihood of observing $y$ given the value of $\varepsilon_L$ and $\gamma$; $g(\varepsilon_L)/g(\gamma)$ define the prior distribution of $\varepsilon_L/\gamma$, which encodes our knowledge of the parameter value before seeing $y$. The denominator of the RHS of Eq. (22) only serves as a normalization factor and is treated as a constant. In practice, the proportionality relationship in Eq. (23) is sufficient for our purpose.

$$p(\varepsilon_L, \gamma|y) \propto f(y|\varepsilon_L, \gamma)g(\varepsilon_L)g(\gamma) \tag{23}$$

Next, we specify the likelihood function and the prior distributions at the RHS of Eq. (23). Given the values of $\varepsilon_L$ and $\gamma$, we can calculate the expected fold change ($\mu$) with main text **Equation 1**. In our experiments, however, several independent replicate measurements of fold change are made to suppress random error, which are expected to be normally distributed about the theoretical value $\mu$. We thus have

$$f(y|\varepsilon_L, \gamma) = \frac{1}{(2\pi\sigma^2)^{N/2}} \prod_{i=1}^{N} exp\left(\frac{-[y_i - \mu(\varepsilon_L, \gamma)]^2}{2\sigma^2}\right) = Normal\left\{\mu(\varepsilon_L, \gamma), \sigma\right\}, \tag{24}$$

where $N$ is the total number of measurements made for $y$. As we've introduced an additional parameter $\sigma$ that describes the width of measurement distribution about the expected value in our statistical model, our complete posterior distribution becomes

$$p(\varepsilon_L, \gamma, \sigma|y) \propto f(y|\varepsilon_L, \gamma, \sigma)g(\varepsilon_L)g(\gamma)g(\sigma), \tag{25}$$

where $g(\sigma)$ is the prior distribution of $\sigma$.

Now with the likelihood function specified, our only task left before having a complete posterior distribution is to define the three prior distributions at the RHS of Eq. (25).

As seen in main text **Equation 1**, $\varepsilon_L$ affects the gene expression level effectively through the composite factor $e^{-\varepsilon_L}/K^2$ in our two-domain model, which has to be evaluated as a whole in the Bayesian inference. Nonetheless, as only 3 of the total 24 mutants investigated in this work contain mutations of direct ligand-binding residues, the change of the inferred composite factor from the WT value for any mutant is attributed solely to the variation of $\varepsilon_L$ for intuitive comparison. However, we note that for the 3 mutants that do contain mutations of ligand-binding residues (P105M, I174K and F177S) ***Leander et al.*** (***2020***), such change could result from the variation of $K$ as well. For example, the triple mutant P176N-I174K-F177S (PIF) has the highest $\varepsilon_L$ among all the queried mutants, which is intuitive considering that it contains mutations of two ligand-contacting residues.

In practice, we set $K$ to 1 nM for convenience and assign a Gaussian prior to $\varepsilon_L$ (Eq. (26)), which admits the ability of point mutations to change intra-domain energetics by a few $k_BT$ while permitting more extreme scenarios ***Chure et al.*** (***2019***); ***Daber et al.*** (***2011***); ***Reichheld et al.*** (***2009***). The apparent binding affinity between our ligand (anhydrotetracycline; aTC) and TetR calculated with $K$ = 1 nM and $\varepsilon_L$ of values within one standard deviation from the mean of g($\varepsilon_L$) well contain those of a range of TetR mutants reported in previous experimental works ***Scholz et al.*** (***2000***); ***Schubert et al.*** (***2004***). Like in the case of $\varepsilon_D$, we focus on the difference between the inferred $\varepsilon_L$ of mutants and WT rather than their absolute value (main text **Figure 3** and **Figure 4—figure Supplement 2**).

$$g(\varepsilon_L) = Normal\{5.5, 2.5\} \tag{26}$$

Similarly, we assign a normal distribution for the inter-domain coupling $\gamma$, which permits a cooperative free energy on the order of a few $k_BT$ ***Motlagh et al.*** (***2014***), and the rare situation where a point mutation can reverse the sign of cooperativity between the two domains ***Scholz et al.*** (***2004***).

$$g(\gamma) = Normal\{5, 2.5\} \tag{27}$$

Lastly, following Chure et al. ***Chure et al.*** (***2019***), the prior distribution of $\sigma$ is given by a half normal distribution (Eq. (28)), with $\phi$=0.05. Such a choice restricts most fold change values generated by the statistical model to stay within the physical bounds of 0 and 1 (see main text **Equation 1**), while permitting rare exceptions that extend slightly beyond the bounds due to experimental noise.

$$g(\sigma) = \sqrt{\frac{2}{\pi\phi^2}} exp\left(\frac{-\sigma^2}{2\phi^2}\right) \tag{28}$$

To check the validity of the chosen prior distributions, we inspect if the simulated induction data based on them comply with our physical understanding of the system in the next step.

### ii. Prior predictive checks

With the statistical model and the prior distributions in hand, we then simulate a set of induction data through such proposed data generation process, and check if the simulated fold changes stay mostly within the physical bounds of 0 and 1.

Specifically, we first draw 1000 sets of leakiness ($\varepsilon_D$), $\varepsilon_L$, $\gamma$ and $\sigma$ values from their prior distributions. For each set of parameters, we then calculate the expected fold change $\mu$ and draw four fold changes from the likelihood function (Eq. (24)) at each one of the 12 ligand concentrations used for experimental induction curve measurements. This matches the number of experimental measurements we made for most mutants. The prior distribution of leakiness is chosen as a half normal distribution centered at 0.005 with a standard deviation of 0.3 (see main text **Figure 3—figure Supplement 2**A), which well covers the range of experimental leakiness we see and expect.

As shown in main text **Figure 3—figure Supplement 2**B, the $5^{th}$ percentile of the simulated fold change measurements has the characteristic shape of an induction curve. $95\%$ of the simulated data falls between $\sim$ 0.05 and 1.1, and the $99^{th}$ percentile extremes are bound between -0.1 and 1.2, which agree with our physical expectation considering the noise in biological measurements.

Satisfied with our prior choices, we go on to check the validity of the complete statistical model and our computational algorithm used for inference in the next step.

### iii. Simulation based calibration

With confidence in our chosen prior distributions, we proceed to check if our complete statistical model and computational algorithm allow for a faithful inference of the parameter values given the corresponding fold change data.

To do so, for each of the 1000 sets of parameters (leakiness, $\varepsilon_L$, $\gamma$ and $\sigma$) drawn for prior predictive check with its simulated data $\tilde{y}$, we estimate its posterior distribution p($\varepsilon_L$, $\gamma$, $\sigma \,|\, \tilde{y}$) given the leakiness and see how well we can recover the true parameter values. The posterior distributions are sampled using Markov chain Monte Carlo (MCMC). Specifically, for each $\tilde{y}$, one million Monte Carlo (MC) sweeps are performed to sample its posterior distribution. In each MC sweep, random moves of $\varepsilon_L$, $\gamma$ and $\sigma$ are proposed sequentially, which are accepted based on the probability specified in Eq. (25) following the Metropolis algorithm. Parameter values after every 1000 MC sweeps are recorded as the posterior samples (see the main text section "*Materials and methods*" for the specific code used for MC sampling).

To assess the validity of our complete statistical model and computational algorithm, the sampled posterior distributions for the 1000 prior predictive draws are then examined using several diagnostic methods.

First, for any Bayesian model, the posterior distribution averaged over prior predictive draws (of large enough sample size) should always recover the prior distribution, as is proven in Eq. (29). Here, $\tilde{\theta}$ is the prior predictive draw of parameter, $\tilde{y}$ is its simulated data and $\theta$ is the inferred parameter. Any deviation between the distribution of inferred $\theta$ and its prior distribution indicates mistakes in either the prior predictive sampling or the estimation of posterior distributions.

$$\pi(\theta) = \iint d\tilde{y} d\tilde{\theta}\ p(\theta|\tilde{y}) f(\tilde{y}|\tilde{\theta}) g(\tilde{\theta})$$
$$= \int d\tilde{y} \frac{f(\tilde{y}|\theta) g(\theta)}{\pi(\tilde{y})} \pi(\tilde{y})$$
$$= g(\theta) \tag{29}$$

As shown in main text **Figure 3—figure Supplement 3**, the average of our inferred posterior distributions (brown) of $\varepsilon_L$, $\gamma$ and $\sigma$, accurately recover the corresponding distributions of the ground truth values (blue) of the prior predictive draws. Therefore, our model well satisfies the self consistency condition.

Second, we performed another ensemble level test of posterior distribution sampling using the rank statistics. Specifically, for each of the 1000 prior predictive draws of parameters $\tilde{\theta}$ ($\theta$=$\varepsilon_L$, $\gamma$ or $\sigma$) and its simulated data $\tilde{y}$, we've collected 1000 MC samples $\theta'_i$ ($i$ ∈[1,1000]) to estimate its posterior distribution $p(\theta'|\tilde{y})$ as detailed above. We then count how many of the posterior samples $\theta'_i$ are larger than the ground truth $\tilde{\theta}$, which is recorded as r($\tilde{\theta}$) (r($\tilde{\theta}$) ∈ [0,1000]). As proved by Talts et al. ***Talts et al.*** (***2018***), if $\theta'_i$ are sampled independently from the correct posterior distribution, the rank statistics (r($\tilde{\theta}$)) of the prior predictive draws should be uniformly distributed (over the integers $\{0, 1, ..., 1000\}$ in our case).

Several visualizations of the rank statistics of our prior predictive draws relative to the corresponding posterior samples are shown in main text **Figure 3—figure Supplement 4**. Both the histograms (main text **Figure 3—figure Supplement 4**A) and the empirical cumulative distribution function (ECDF) plots (main text **Figure 3—figure Supplement 4**B and C) show that the rank statistics of $\varepsilon_L$, $\gamma$ and $\sigma$ are all uniformly distributed. While the histogram provides a general and interpretable way of checking uniformity, the ECDF is more sensitive to small deviations especially at small and large ranks. The green bands in main text **Figure 3—figure Supplement 4**A-C show the $99^{th}$ percentile expected from a true uniform distribution. The ECDF difference in main text **Figure 3—figure Supplement 4**C is obtained by subtracting the theoretical cumulative distribution function of a uniform distribution from the observed ECDF, which makes deviations more evident if existent.

Third, besides examining the ensemble averaged behavior of the inferred posterior distributions, we further compute the posterior z-score and posterior contraction of each posterior distribution to see how well it recovers the true values of the corresponding parameters. As stated above, for each of the 1000 prior predictive draws of parameters $\tilde{\theta}$ ($\theta$=$\varepsilon_L$, $\gamma$ or $\sigma$) and its simulated data $\tilde{y}$, we've estimated its posterior distribution $p(\theta'|\tilde{y})$ using MCMC. The posterior z-score is defined as

$$z = \frac{M[p(\theta'|\tilde{y})] - \tilde{\theta}}{\sqrt{V[p(\theta'|\tilde{y})]}} \tag{30}$$

where $M$ and $V$ denote mean and variance respectively. It quantifies how accurately the posterior recovers the true value of the inferred parameter. Apparently, a smaller/larger (absolute values of) z-score indicates that the posterior is concentrated around/away from the true parameter value. However, as posterior z-score reports the relative magnitude of the bias and width of the posterior distribution, it doesn't reflect the precision of the inference.

Another characterization of the posterior of a parameter, known as posterior contraction, is defined as the ratio of the posterior variance to the prior variance subtracted from one (Eq. (30)). Posterior contraction quantifies to what degree the data inform the posterior. Posterior contraction near 0 indicates that the posterior inference is largely influenced by the prior distributions, and is poorly informed by the data; while posterior contraction close to one indicates that the data are much more informative than the prior distributions.

$$c = 1 - \frac{V[p(\theta'|\tilde{y})]}{V[g(\theta)]} \quad (31)$$

The z-score and contraction analysis of posterior distributions can identify a series of pathologies of the statistical model ***Schad et al.*** (***2020***). Ideally, a posterior distribution should stay close to [1,0] on the plane spanned by the axis of contraction and z-score, which means that the posterior accurately recovers the true parameter value with high precision.

The posterior z-score and contraction of 970 of our prior predictive draws are shown in main text **Figure 3—figure Supplement 5**A, where z-scores of all parameters are clustered around 0, indicating high accuracy of their inferences. On the other hand, while the contractions of $\varepsilon_L$ and $\sigma$ are close to 1, the contraction of $\gamma$ is tailed with a minimum around 0.4 and a median of 0.972. Although the contraction of $\gamma$ is not as ideal as those of $\varepsilon_L$ and $\sigma$, it still reflects that the inference of $\gamma$ is reasonably well informed by the data ***Chure et al.*** (***2019***); ***Schad et al.*** (***2020***).

In the above analysis, we excluded 30 prior predictive samples satisfying the criteria given in Eq. (32), which essentially correspond to flat induction curve. The parameter values of these samples cannot be faithfully inferred due to the model degeneracy of flat induction curves as discussed in section 2 and main text **Figure 2—figure Supplement 3**. As the smallest $seperation$ (Eq. (32)) of our experimental induction data is 8.6, (except that of G102D, see main text **Figure 2—figure Supplement 3**), we don't encounter such problem in the inference using real data.

$$seperation = \left| \frac{\mu(c = 1000\ nM) - \mu(c = 0)}{\sigma} \right| < 2 \quad (32)$$

It's noted that the contraction of $\gamma$, while reasonable, is not as ideal as those of other parameters (main text **Figure 3—figure Supplement 5**A), which prompts a physical explanation. Accordingly, a closer examination of our physical model reveals the intrinsic difficulty of inferring $\gamma$ under two scenarios.

First, when the sum of $\gamma$ and $\varepsilon_L$ is large to the extent that the condition of Eq. (33) is satisfied, the main text **Equation 1** is reduced to Eq. (34). For example, when the left-hand side (LHS) of Eq. (33) is smaller than 0.05, the corresponding sum of $\varepsilon_L$ and $\gamma$ will be greater than 16.8 $k_B T$.

$$e^{-\varepsilon_L - \gamma} \left(\frac{c}{K}\right)^2 \ll 1,\ c = c^{max} = 1000\ nM \quad (33)$$

$$FC = \left( 1 + \frac{R^* e^{-\varepsilon_D}}{1 + e^{-\varepsilon_L} \left(\frac{c}{K}\right)^2} \right)^{-1} \quad (34)$$

Under such scenario, the induction curve will be insensitive to the value of $\gamma$, as long as it's large enough so that Eq. (33) holds. This reversely causes the wider posterior distribution of $\gamma$ when it is inferred from such induction data, leading to its lower contraction compared with other parameters (Eq. (31)). On the other hand, Eq. (34) shows that its corresponding induction curve will either saturate at $FC$=1, or has no saturation, featuring a sharply varying tail. We note that an induction curve with a sharply varying tail region may arise when Eq. (33) is not satisfied, under those conditions parameter inference does not suffer any difficulty.

Second, highly precise inference of $\gamma$ will also be difficult when Eq. (35) is true. Likewise, under such condition, main text **Equation 1** is effectively reduced to Eq. (34), except that the corresponding induction curve is required to saturate at $FC$=1.

$$\begin{cases} e^{-\gamma} \ll 1, & e^{-\varepsilon_L} \left(\frac{c}{K}\right)^2 \gg R^* e^{-\varepsilon_D} \\ e^{-\varepsilon_L - \gamma} \left(\frac{c}{K}\right)^2 \ll 1, & \text{smaller c than required above.} \end{cases} \quad (35)$$

Indeed, when we exclude the prior predictive draws with $\mu$($c$=1000 nM)>0.97 from the analysis, the average inferential performance of the model improved noticeably especially in the case of $\gamma$, whose new median contraction is above 0.997 (main text **Figure 3—figure Supplement 5**B). The remaining few instances of lower $\gamma$ contraction still corresponds to large sums of $\gamma$ and $\epsilon_L$, however, with the induction curve featuring a sharply varying tail region, as discussed above.

In our experimental dataset, such inference difficulty is only observed in the case of C203V, Y132A-C203V and C203V-G102D-L146A due to their large $\gamma$ and $\gamma + \epsilon_L$ values (see main text **Figure 3**, **Figure 3—figure Supplement 10** and **Figure 4**). As shown in main text **Figure 3—figure Supplement 10**, the inference results for the other 20 mutants stay highly precise and virtually unchanged after increasing the standard deviation of the Gaussian prior of $\gamma$ ($g_\gamma^{std}$) from 2.5 to 5 $k_B T$. This demonstrates that the inference results for these mutants are strongly informed by the induction data and there is no difficulty in the precise inference of the parameter values. On the other hand, the inferred $\gamma$ values (especially the upper bound of the 95% credible region) for C203V, Y132A-C203V and C203V-G102D-L146A increased with $g_\gamma^{std}$. This is because the induction curves in these cases are not sensitive to the value of $\gamma$ given that it's large enough as discussed above. Hence, when unphysically large $\gamma$ values are permitted by the prior distribution, they could enter the posterior distribution as well.

Such difficulty in the precise inference of $\gamma$ values for these three mutants however, doesn't compromise the ability of our model in accurately capturing the comprehensive set of induction data (see part iv below). Additionally, the increase of the inferred $\gamma$ value of C203V at the use of larger $g_\gamma^{std}$ complies with the results presented in main text **Figure 4**, which show that the effect of C203V on $\gamma$ tends to be compromised when combined with mutations closer to the domain interface.

Now with confidence in our statistical model and its computational implementation, as well as understanding of the inferential limitations intrinsic to the biophysical model, we proceed to see how well they capture the experimental data.

#### iv. Posterior predictive checks

We now apply our statistical model to the inference of the biophysical parameters of our experimentally tested mutants and inspect how well it captures the corresponding induction data. The same parameter estimation procedure is applied to all the 24 mutants tested except G102D (main text **Figure 3** and **Figure 4—figure Supplement 2**), and we describe the case of G102D-Y42M-I57N here as an example.

First, the $\epsilon_D$ of G102D-Y42M-I57N is calculated to be $1.63^{+0.01}_{-0.02}\ k_B T$ (main text **Figure 3**) based on its leakiness measurements of 0.0423±0.0005 (mean±SEM) and Eq. (21) (the $\epsilon_D$ of WT is taken as 0 which has a leakiness of 0.0086). Next, with the average leakiness and the full induction data $y$, we generated 1000 sets of parameters $\{\epsilon_L, \gamma, \sigma\}$ using MC sampling as estimation of the posterior distribution $p\left(\epsilon_L, \gamma, \sigma | y\right)$. The reported inference result of an individual parameter $\theta$ is the median of the marginalized posterior $p(\theta|y)$ with the error bars showing the $95\%$ credible interval (main text **Figure 3** and **Figure 4—figure Supplement 2**).

To see how well the statistical model and the posterior parameters capture the experimental data from which they are inferred, we generated induction data using each of the 1000 posterior samples (like we did for the prior predictive draws) and inspect how well it recapitulates the experimental observation. The tight joint and marginal distributions in main text **Figure 3—figure Supplement 6**A show that the parameters are inferred with high precision. A moderate correlation is observed between $\epsilon_L$ and $\gamma$, while they are more symmetric to $\sigma$. More importantly, the simulated induction curves using the posterior samples follow the experimental measurements closely, which all fall within the $99^{th}$ percentile. Thus, our statistical model accurately describes the experimental observation, which is also shown in the posterior predictive checks of the other mutants (main text **Figure 3** and **Figure 4—figure Supplement 2**).

### 4. Prediction of mutation combinations based on the additive model

After characterization of the 15 mutants shown in main text **Figure 3**, we can generate prediction of combined mutants assuming additivity. Specifically, the biophysical parameters of a mutant containing two characterized mutations (mut1 and mut2) is predicted based on main text **Equation 4** and **5**. The two equations are first used to calculate the $\epsilon_D$ of the combined mutant, and then applied to every one of the 1000×1000 pairs of the posterior samples ($\epsilon_L$ and $\gamma$) of mut1 and mut2. We thus get the $\epsilon_D$ value with one million sets of $\{\epsilon_L, \gamma\}$ as estimation of the biophysical parameters of the combined mutant. The medians of the marginal distributions of $\epsilon_L$ and $\gamma$ are then reported as the predicted parameter values for the combined mutant, where the error bars show the $95^{th}$ percentile (main text **Figure 4—figure Supplement 2**). In the basic additive model, we set $\alpha_{1,p} = \alpha_{2,p} = 1$ in main text **Equation 4**, where $p$ represents any one of $\epsilon_D$, $\epsilon_L$ and $\gamma$ (**Figure 4—figure Supplement 1-2**). Systematic deviations between the experimental data of combined mutants and additive model prediction should manifest how important is epistasis between the mutations. This is rationalized in the modified additive model, where the values of $\alpha_{1,p}$ and $\alpha_{2,p}$ are adjusted to account for epistasis (main text **Figure 4**).

To assess the success of prediction, we compare the experimental induction curve with that generated using the parameters predicted by the (adjusted) additive model. Specifically, we calculate the theoretical fold change at the 12 ligand concentrations using the calculated $\epsilon_D$ and each set of $\epsilon_L$ and $\gamma$ with main text **Equation 1**. Here, $R^*$ is calculated to be 115 based on the WT leakiness, whose $\epsilon_D$ is set to 0. The $95^{th}$ percentile of thus generated one million induction curves are compared with the experimental induction data to assess the quality of prediction (main text **Figure 4** and **Figure 4—figure Supplement 1**).

### 5. Calculation of the apparent binding affinities to ligand and operator

The apparent biding affinity of TetR to the ligand (aTC) and operator ($tetO2$) used in our experiments are calculated as those of the $L_I D_I$ conformation of the repressor for it has the lowest free energy among all unbound states. Accordingly, the apparent dissociation constant of WT TetR to ligand is calculated as

$$K^{WT}_{apparent} = \sqrt{e^{\epsilon^{WT}_L}} \cdot 1 = 27.1\ (nM), \tag{36}$$

which is converted to a binding free energy of -17.4 $k_B T$, close to the reported experimental value -16.8 $k_B T$ ***Schubert et al.*** (***2004***).

Similarly, according to Eq. (18), the apparent operator binding affinity of WT TetR is estimated as

$$\epsilon_{R_A O} + \epsilon_D = -ln\left(\frac{\frac{1}{leakiness} - 1}{[R_{tot}]}\right) \sim -16.4\ (k_B T), \tag{37}$$

where $[R_{tot}]$ is evaluated based on the repressor copy number per cell of 5000 and the E. Coli cell volume of 1 $\mu m^3$ ***Kubitschek and Friske*** (***1986***). This is comparable to the reported experimental value of -18.8 $k_B T$ ***Kędracka-Krok and Wasylewski*** (***1999***). However, it's noted that without direct measurement of the repressor concentration in the cell, this is only meant to be an order of magnitude comparison.

### 6. Epistasis between C203V, Y132A and other mutations

To probe the epistasis between mutations in TetR, we seek systematic deviations between the additive model prediction and experimental data of combined mutants. For example, our results indicate that quenching C203V's effect on $\epsilon_D$ is a common modification of the additive model that improves its agreement with the experiments (main text **Figure 4** and **Figure 4—figure Supplement 2**). The only exception is the mutant Y132A-C203V, where the additive model prediction on $\epsilon_D$

is already close. However, we note that Y132A is the most distant mutation from the DNA-binding residues among all that are paired with C203V (main text **Figure 3—figure Supplement 1**), residing above the ligand as C203V does. Thus, it falls in line with the reasoning of the epistasis in other combined mutants containing C203V (main text **Figure 4**).

On the other hand, despite its proximity to the ligand-binding residues, we find that Y132A's effect on $\varepsilon_L$ tend to be compromised when combined with other mutants, as is strikingly reflected in Y132A-C203V (main text **Figure 4** and **Figure 4—figure Supplement 2**). This is indeed a commonly observed trend in the epistasis of other Y132A containing combined mutants. Tuning down Y132A's contribution to $\varepsilon_L$ in all cases leads to better additive model prediction, while tuning down that of the other mutation (R49G and PIF) resulted in worse performance (main text **Figure 4** and **Figure 4—figure Supplement 2**). Similar to the case of C203V (discussed in the main text), the common epistatic interactions observed for Y132A here offers a possible explanation for why it doesn't rescue a range of dead mutations despite being able to enhance the allosteric response of TetR by itself ***Leander et al.*** (***2020***).

### 7. Mutation selection for two-domain model analysis

In this work, there are 24 mutants studied in total including the WT, and they contain mutations at 21 WT residues. We did not perform model parameter inference for the mutant G102D because of its flat induction curve (see the second subsection of section 2 and main text **Figure 2—figure Supplement 3**). Therefore, there are 23 mutants analyzed in main text **Figure 5**.

Measuring the induction curve of a mutant involves a significant amount of experimental effort, which therefore is hard to be extended to a large number of mutants. Nonetheless, we aim to compose a set of comprehensive induction data here for validating our two-domain model for TetR allostery. To this end, we picked 15 individual mutants in the first round of induction curve measurements, which contains mutations spanning different regions in the sequence and structure of TetR (main text **Figure 3—figure Supplement 1**). Such broad distribution of mutations across LBD, DBD and the domain interface could potentially lead to diverse induction curve shapes and mutant phenotypes for validating the two-domain model. Indeed, as discussed in the main text section "Extensive induction curves fitting of TetR mutants", the diverse effects on induction curve from mutations perturbing different allosteric parameters predicted by the model, are successfully observed in these 15 experimental induction curves. Additionally, 5 of the 15 mutants contain a dead-rescue mutation pair, which helps us validate the model prediction that a dead mutation could be rescued by rescuing mutations that perturb the allosteric parameters in various ways.

Eight mutation combinations were chosen for the second round of induction curve measurement for studying epistasis, where we paired up C203V and Y132A with mutations from different regions of the TetR structure. Such choice is largely based on two considerations. 1. As both C203V and Y132A greatly enhance the allosteric response of TetR, we want to probe why they cannot rescue a range of dead mutations as observed previously ***Leander et al.*** (***2020***). 2. C203V and Y132A are the only two mutants that show enhanced allosteric response in the first round of analysis. Combining detrimental mutations of allostery in a combined mutant could potentially lead to near flat induction curve, which is less useful for inference ((see the second subsection of section 2).

### 8. The simplicity of the two-domain model

In this work, we aim to build a minimalist model for two-domain allostery with only the most essential parameters for capturing experimental data. Hence, the homodimeric nature of TetR is deliberately ignored here for the simplicity of the two-domain model, which helps promote its mechanistic clarity and potential transferability to other allosteric systems.

Moreover, fewer parameters are needed in a simpler model. Our two-domain model currently uses only three biophysical parameters, which are all demonstrated to have distinct influences on the induction curve (see the main text section "System-level ramifications of the two-domain model”). This enables the inference of parameters with high precision for the mutants, and the quantification of the most essential mechanistic effects of their mutations, provided that the model is shown to accurately recapitulate the comprehensive dataset (main text **Figure 3** and **Figure 4—figure Supplement 2**). Thus, we found it was unnecessary to add another parameter for explicitly describing inter-chain coupling, which would likely incur uncertainty in the inference of parameters due to the redundancy of their effects on induction data, and prevent the model from making faithful predictions.

From a more biological point of view, TetR is an obligate dimer, meaning that the two chains must synchronize for function, supporting the two-domain simplification of TetR for binding concerns. Additionally, as shown in the subsection “Inclusion of single-ligand-bound state of repressor” of section 1, incorporating the dimeric nature of TetR in our model by allowing partial ligand binding does not change the functional form of main text equation 1 in any practical sense.

In summary, we think that the value of a simple physical model is two-fold (e.g., the paradigm Ising model in statistical physics and the classic MWC model), first, its mechanistic clarity and potential transferability makes it a useful conceptual framework for understanding complex systems and establishing universal rules by comparing seemingly unrelated phenomena; second, it provides useful insights and design principles of specific systems if it can quantitatively capture the corresponding experimental data. Thus, given the current experimental data set, we believe it is justified to keep the two-domain model in its current form, while additional experimental data could necessitate a more complex model for TetR allostery in the future.
